# Computational Inference Software for Tetrad Assembly from Randomly Arrayed Yeast Colonies

**DOI:** 10.1101/574376

**Authors:** Nikita A. Sakhanenko, Gareth A. Cromie, Aimée M. Dudley, David J. Galas

## Abstract

Here, we describe an information-theory-based method and associated software for computationally identifying sister spores derived from the same meiotic tetrad. The method exploits specific DNA sequence features of tetrads that result from meiotic centromere and allele segregation patterns. Because the method uses only the genomic sequence, it alleviates the need for tetrad-specific barcodes or other genetic modifications to the strains. Using this method, strains derived from randomly arrayed spores can be efficiently grouped back into tetrads.

## Introduction

In many eukaryotes, including the genetically tractable yeasts *Saccharomyces cerevisiae* and *Schizosaccharomyces pombe*, the filamentous fungus *Neurospora crassa*, and the unicellular green alga *Chlamydomonas reinhardtii*, it is possible to recover all four of the haploid products of a single meiosis, tetrads. These tetrads can be characterized genetically and phenotypically. Tetrad analysis is a powerful technique that is routinely used to make associations between genetic variation and phenotype, uncover gene-gene interactions, and identify non-reciprocal meiotic recombination events (e.g. gene conversions).

The manual processes of isolating, disrupting, and arraying spores in conventional tetrad analysis have limited its application to relatively small-scale studies. The conventional method has two steps that are difficult to automate, isolating tetrads away from unsporulated cells in the culture and capturing the sister spore relationships of the resulting progeny strains by arraying the spores in a gridded pattern. We previously described a method, BEST (Barcode Enabled Sorting of Tetrads) (Ludlow et al. 2013; Scott et al. 2014), that uses a meiosis-specific GFP fusion protein to isolate tetrads by fluorescence-activated cell sorting of tetrads and molecular barcodes to identify sister spores of the same tetrad by DNA sequencing.

Although plasmid-borne tetrad-specific molecular barcodes are well suited for laboratory strains of *S. cerevisiae*, they may not be as useful for organisms that are less genetically tractable (e.g. with low transformation efficiency or poor maintenance of extrachromosomal plasmids) or for the construction of non-genetically modified strains. The problem of reconstructing tetrad information from a large set of randomly arrayed spores can be viewed as two sub-problems: (1) finding a reliable measure that identifies groups of four sister spores (a tetrad); and (2) defining an appropriate search strategy to efficiently traverse a very large set of possible spore groupings while applying this measure. Here, we describe an information-theory-based metric that solves the first sub-problem and software implementing a search strategy utilizing this metric that solves the second. Because the method uses only the genomic DNA sequence of the meiotic products, it can be applied to strains or organisms for which genetic manipulation is difficult or undesirable.

## Results

### The mechanisms of recombination and chromosome segregation produce unique genotypic signatures for each meiosis

Meiosis is a process in which a diploid cell undergoes one round of DNA replication followed by two rounds of chromosome segregation and cell division to produce four recombinant haploid progeny. In the first meiotic division, the two homologs of each chromosome recombine and then segregate to the opposite poles of the meiotic spindle. In the second meiotic division, the two chromatids of each recombinant chromosome segregate with no further recombination (Figure 1A). Therefore, in the absence of rare gene-conversion events, at each position that is heterozygous in the original diploid, exactly two spores will inherit allele “A” and exactly two will inherit allele “B” (Figure 1A). Additionally, because sister chromatids only segregate from each other at the second meiotic division, two of the spores will have identical centromeric alleles for every chromosome, while the other two spores will both have the mirror image of this pattern (Figure 1B). Thus, centromere allele segregation patterns, constrained allele frequencies, and patterns of recombination breakpoints can be viewed as genotypic signatures for individual meioses. These signatures could, in principle, be used to reconstruct tetrads computationally from the DNA sequences of randomly arrayed spores.

**Figure 1.**
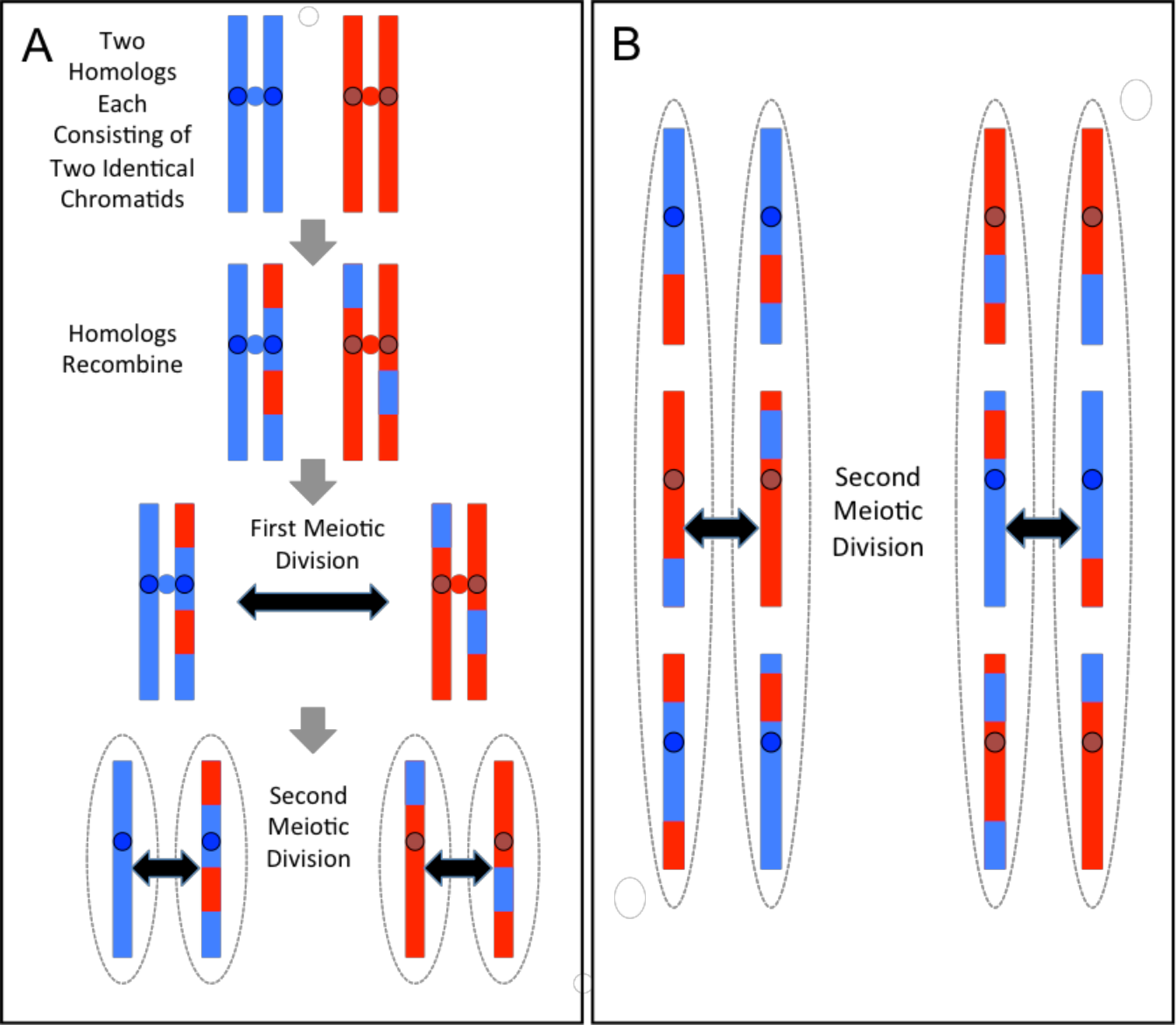
(A) Behavior of a single chromosome during meiosis. In the initial heterozygous diploid (top) there are two copies of the “A” haplotype (blue chromatids) and two copies of the “B” haplotypes (red chromatids). Centromeres are shown as circles. The two “A” centromeres remain paired until the second meiotic division, as do the two “B” centromeres. Spores (haploid meiotic products) are shown as dotted ovals. (B) Segregation pattern shown for 3 chromosomes. For each chromosome, segregation of the “red” or “blue” homologs to the left or to the right side at the first meiotic division occurs at random, but for each chromosome the two sister “red” and the two sister “blue” centromeres always remain paired until the second meiotic division. Therefore, at each centromere the two leftward spores always share the same allele and the two rightward spores share the other allele.

Here, we report a computational method, hereafter “tetrad reconstruction”, that uses the pattern of centromere segregation and the constrained allele frequencies within a tetrad to infer the original sister spore relationships of recombinant progeny. Our method takes genotype data for all of the progeny strains as input and then proceeds in two steps. First, the centromere segregation pattern is used to reduce the number of potential spore patterns to be searched. Then, the constrained allele segregation patterns are used as the signal to identify members of the same tetrad.

### Using centromere segregation patterns to reduce the search space for spores from the same tetrad

With an appropriate metric based on allele frequencies or recombination breakpoints, it is possible to distinguish a group of spores from the same tetrad from other groups of spores. However, applying this metric in an exhaustive, brute force search across all possible groups of spores in a large dataset is computationally demanding. To reduce this search space and simplify the computational problem, we implemented an efficient heuristic based on the segregation patterns of centromeres in tetrads, i.e. the fact that two spores of a tetrad harbor the same alleles at each centromere and the other two spores both share the opposite pattern (Figure 1B). Our heuristic search leverages this property by first attempting to partition the set of all spores into clusters of spores whose centromere-flanking markers are either a perfect match or a complete mismatch. Unless a polymorphic marker is present at the centromere itself, it is not possible to determine the haplotype origin (“A” or “B”) of centromeres with absolute certainty. Therefore, we compute the probability of each centromere coming from haplotype “A” or haplotype “B” based on the alleles of the flanking markers and the empirically-estimated recombination frequency between them and the centromere (**Appendix G**). Given these probabilities, we derive a similarity coefficient between all spores (**Appendix B**) and use a fast greedy algorithm (Clauset 2004) to cluster spores based on these similarity coefficients.

There are *2*^*N*^ unique segregation patterns, where *N* is the number of chromosomes. However, only half of these patterns (*2*^*N-1*^) can uniquely identify a tetrad, since each tetrad contains two patterns opposite to one another (see Figure 1B). Therefore, for organisms with a sufficiently large number of chromosomes, our clustering algorithm based on the centromere segregation heuristic should produce small clusters that contain all members of a given tetrad. For example, in *S. cerevisiae*, which has sixteen chromosomes, the chance of two tetrads sharing a centromere segregation pattern is 1/2^15^. However, in an organism with fewer chromosomes, such as *S. pombe*, which has only three, more than one tetrad will often be assigned to a single cluster. In addition, factors such as sequencing errors, or crossovers between a centromere and its flanking markers, can also lead to false positive or false negative tetrad assignments. Therefore, although it reduces the search space for subsequent steps, this heuristic alone is not sufficient to accurately reconstruct tetrads.

### Using information theory to reconstruct tetrad relationships based on 2:2 allele segregation

The metric that we chose to use for unambiguously identifying members of the same tetrad is based on the fact that at each marker locus in a tetrad, two spores inherit the “A” allele and two spores inherit the “B” allele (Figure 1). Thus, the allele patterns within a tetrad are constrained and knowledge of the genotype of one spore changes the allele probabilities for the other three spores. For example, at every position where an “A” allele is observed in one spore, the probability of the “A” allele in any of the remaining three spores of the same tetrad changes from 50% to 33%. In contrast, knowledge of the genotype of a spore from one tetrad does not affect the allele probabilities in spores from different tetrads. As such, tetrad-specific relationships can be viewed as dependencies among the four allele vectors of a tetrad (one vector for each spore genotype), and such dependencies can be detected using methods from information theory. In contrast to the centromere heuristic, this constrained allele frequency approach uses a much larger number of genotyped markers, making the approach less sensitive to individual genotyping errors and more successful at disambiguating tetrad assignments.

Mutual information is a well-known measure that quantifies the amount of dependency between two categorical variables (**Appendix A**), and interaction information (McGill, 1954) is a multivariable generalization of this measure (**Appendix A**). While interaction information has a number of drawbacks (Bell, 2003; Jakulin and Bratko, 2004; Sakhanenko and Galas, 2011), it can be used in principle to devise measures of dependency among any number of variables. Because the genotypes of a group of spores from the same tetrad are a set of dependent variables, they should produce a strong interaction information signal. In contrast, if the genotype of a spore from a different tetrad (an independent variable) is then added to the group, the interaction information should be close to zero. Thus, an interaction information approach might be able to identify groups of four strains that were sister spores from the same tetrad. Furthermore, in cases where spores are missing or inviable, the allele vectors of the remaining spores will still show dependencies among one another, detectable by mutual information between two spores or interaction information between three spores.

To test the ability of interaction information to identify groups of spores from the same tetrad, we generated a simulated dataset of 100 *S. cerevisiae* tetrads with 1000 markers, 1% noise (genotyping errors), and 5% missing data (**Appendix F**). We then calculated interaction information for the real tetrads and for groups of 4-spores derived from different tetrads. On average, true tetrads scored highly by this metric, while incorrectly grouped spores had scores centered on zero. However, while most incorrect groups of spores scored poorly, there were a number of false positives, i.e. incorrect groupings that scored as highly as some of the true tetrad groups (Figure 2A). A similar result was seen with groups of three spores from the same tetrad versus incorrect groupings of 3-spores, and the overlap was even stronger in this case (Figure 2B). These results suggest that noise in the data limits the ability of interaction information to cleanly distinguish correct groups of three or four spores derived from the same tetrad from incorrect groupings.

**Figure 2.**
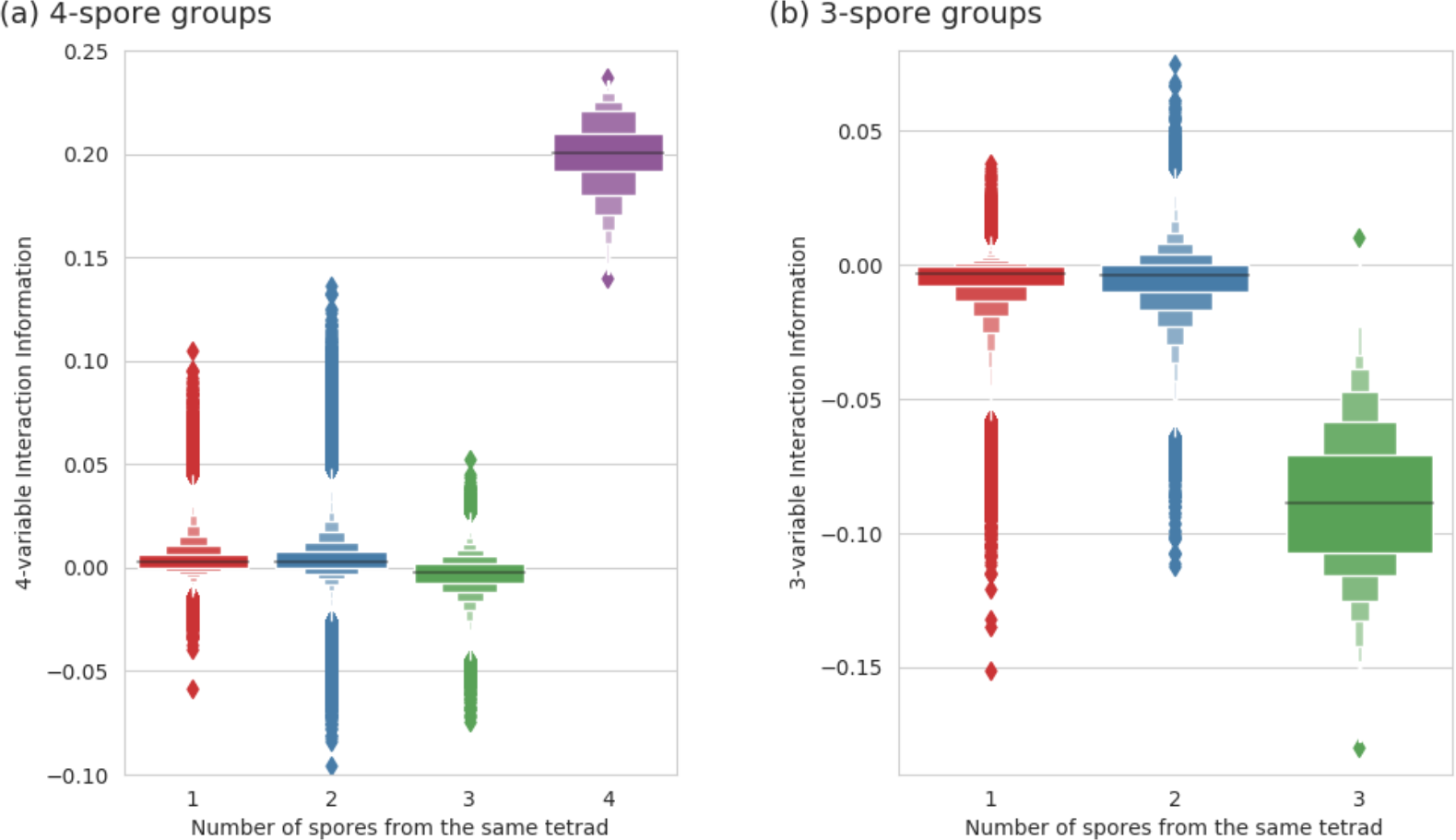
Interaction Information computed on groups of (a) 4 spores and (b) 3 spores. All measures were computed on the simulated *S. cerevisiae* data (1000 markers, 100 tetrads, 1% noise, and 5% missing data). Panel (a) distinguishes four possible categories of 4-spore groups based on the number of spores derived from the same tetrad (shown on the x-axis) and shows a letter-value (LV) plot for each category. The purple LV plot shows the Interaction Information scores for all possible real tetrads. Similarly, panel (b) shows the scores for three different categories of 3-spore groups. For the categories where all spores are from different tetrads (the red LV plots in (a) and (b)), only 2 million randomly sampled groups are shown. Note that the Interaction Information signal associated with true sister spores is negative at the 3-spore level and positive at the 4-spore level, while noise is clustered around zero in both cases.

To explore this problem further, we considered the behavior of interaction information at multiple levels of complexity. As expected, a real tetrad has strong interaction information signal at the 4-spore level as well as at the 3-spore level for all subgroups of three spores (Figure 3a and S7). In contrast, when an incorrectly assembled tetrad has a relatively high interaction information signal at the 4-spore level due to noise, this signal does not extend to its 3-spore subgroups (Figure 3a, green oval), a result that suggests that noise at the 3- spore and 4-spore levels is not correlated. We also observed similar, but noisier, patterns between the 2- and 3-spore levels (Figure 3b, green oval). Thus, we hypothesized that combining interaction information at the 4-spore and 3-spore levels or from the 3-spore and 2-spore levels should substantially strengthen the signal separating real 4- and 3-spore tetrads from false ones.

**Figure 3.**
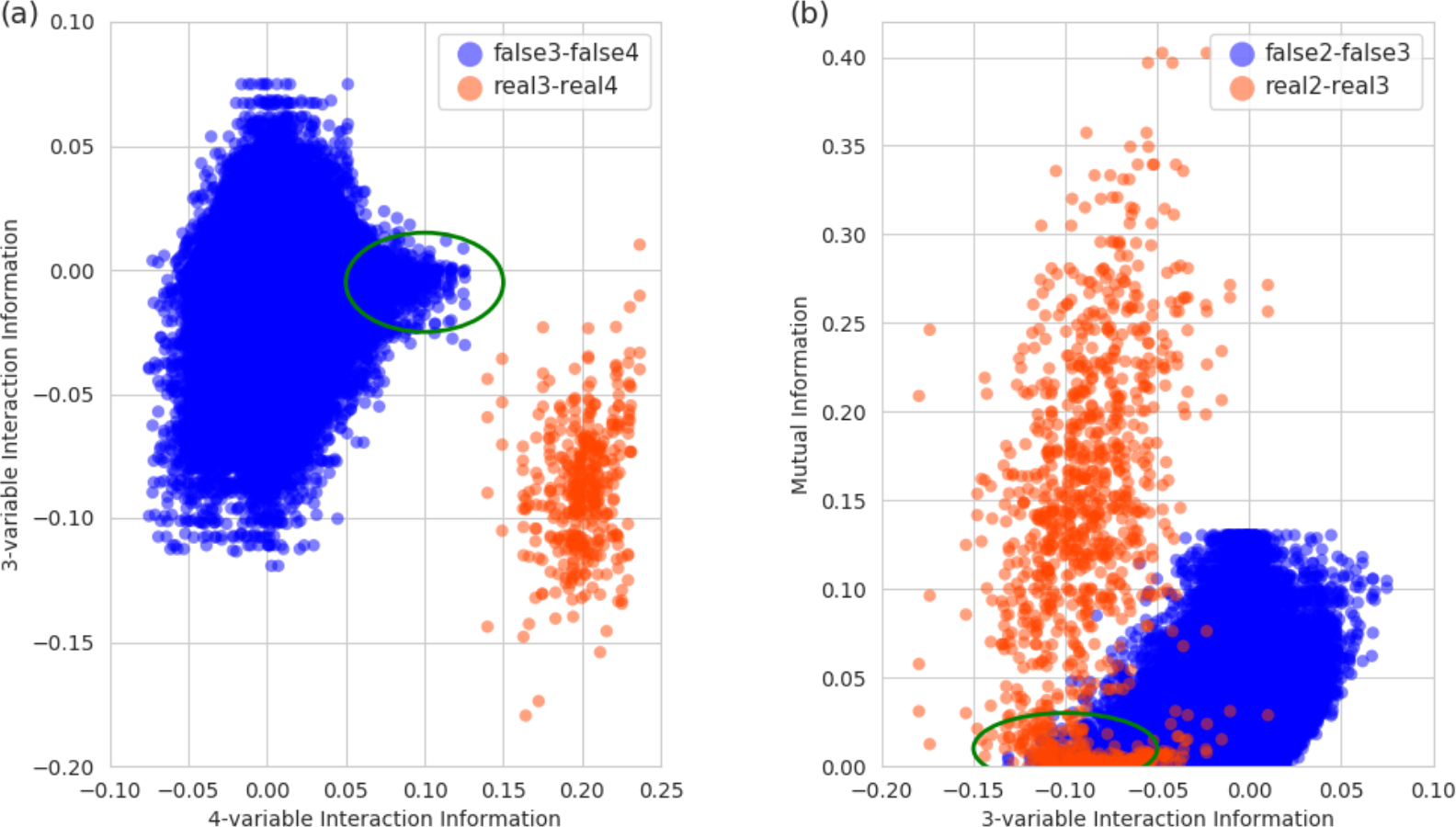
Comparison of the amount of information between (a) 2-spore and 3-spore levels and between (b) 3-spore and 4-spore levels as measured by interaction information. All measures were computed on the same data as in Figure 2. Panel (a) shows the scatter plot of scores computed on various groups of 4 spores and their 3-spore subsets while panel (b) shows the scores for groups of 3 spores and their 2-spore subsets. Each group is colored red if all spores of the group came from the same tetrad and blue if at least two spores came from a different tetrad. The scores of the red sets are plotted in their entirety, whereas for the blue sets we randomly selected 2 million groups. Green ovals indicate the situation when, due to noise, the interaction information shows relatively high signal at one level for a group of spores derived from more than one tetrad. Note that at the level below, the signal from these groups is close to zero.

To combine interaction information at different spore-number levels, we used a measure based on differential interaction information, called “delta” (Galas et al., 2014; Sakhanenko and Galas, 2015; Galas and Sakhanenko, 2016). Differential interaction information quantifies the change in interaction information that occurs when a new variable is added to a set of existing variables. Unlike the interaction information measure, differential interaction information is not symmetric, but is specific to which variable is added. A symmetric measure results from the product of differential interaction information with all possible choices of the added target variable, and this product is “delta” (**Appendix A**).

Consistent with our hypothesis, delta performed significantly better than interaction information in distinguishing groups of spores from the same tetrad from incorrect groupings (see Figure 4, also Figures S6 and S7). By combining information at different degree levels, the delta measure allowed us to distinguish the real tetrads from all other 4- spore groups, with a false positive rate of zero using the test dataset. Furthermore, the difference between real tetrads and incorrect 4-spore groupings was orders of magnitude larger when measured with delta as opposed to interaction information. This difference can also be seen at the 3-spore level: groups of 3 sister spores are separated away from other 3-spore groups if we use delta (see Figure 4d) as opposed to interaction information (see Figure 4c) where these two distributions of 3-spore groups are more ambiguous.

**Figure 4.**
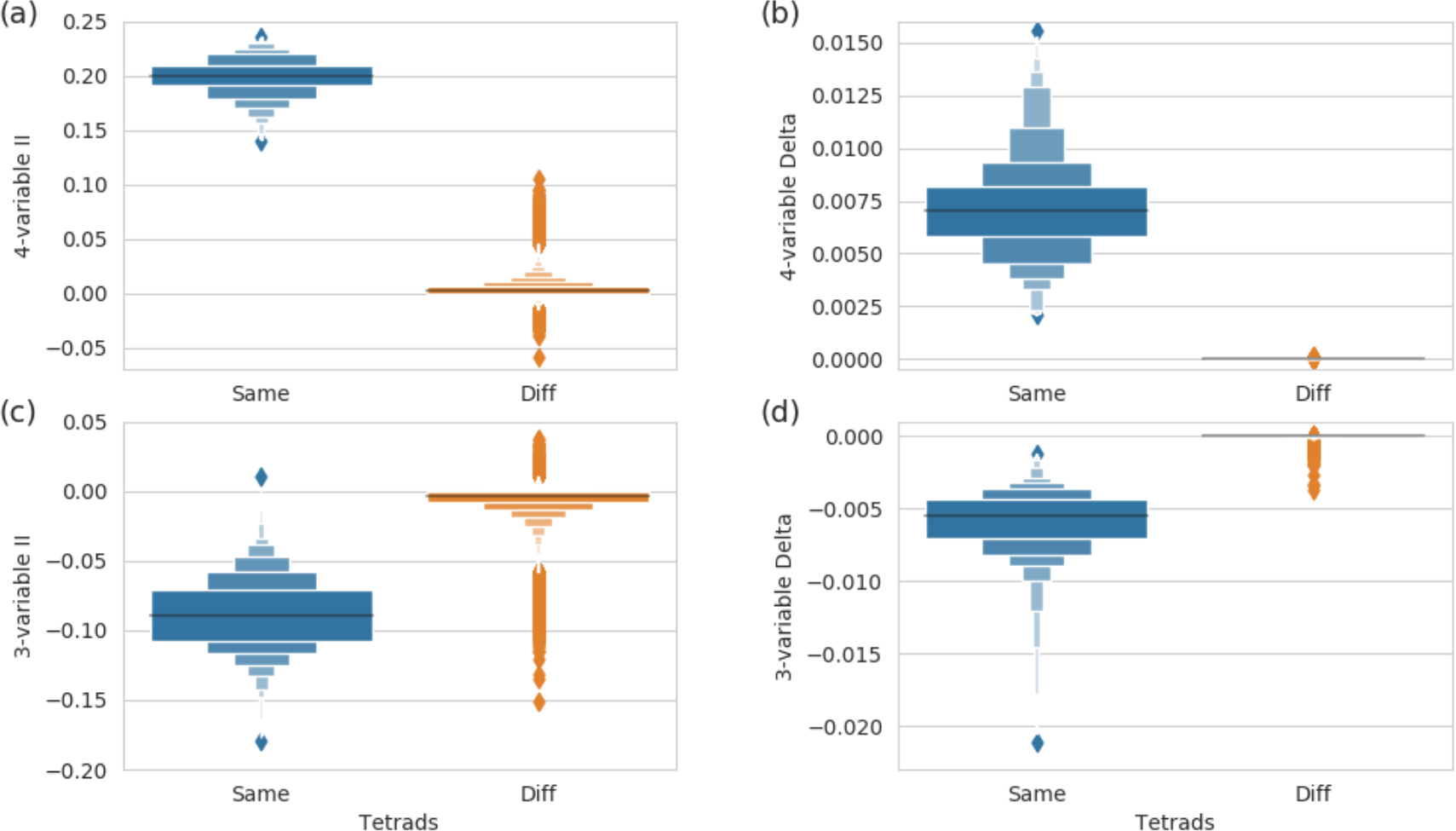
Comparison of the ability of Interaction Information and the delta measure to distinguish groups of sister spores from all other groups. Panel (a) shows LV plots of Interaction Information scores computed on groups of 4 spores from the same tetrad (in blue) and on all other groups of 4 spores (in green). Panel (b) shows delta scores computed on the same 4-spore sets. Similarly, panels (c) and (d) show Interaction Information and delta scores computed on triplets of sister spores (in blue) and triplets of spores from different tetrads (in green). All measures were computed on the same data as in Figure 2.

### Thresholds for identifying spores from the same tetrad and validation using 2:2 allele segregation

To construct a classifier that uses delta for tetrad reconstruction, we need to identify a threshold that distinguishes true-positive tetrads from false-positive groups of spores with high likelihood. We establish this threshold by randomizing the dataset that will be analyzed in order to produce a null distribution consisting of mostly non-tetrad groups. Computing delta scores for all the elements of the null distribution, the delta threshold is then defined based on a user-defined *p*-value. The default for 4-spore tetrads is 0.05, but our software allows the user to adjust this parameter. A group, whose delta score is above the threshold, is then identified as a candidate tetrad for validation. We note that the null distributions obtained by this permutation also include a small proportion of groups of spores from the same tetrad, so the cutoffs are, in practice, slightly conservative.

In a real tetrad with no gene conversions, every marker segregates 2:2, i.e. two spores inherit the “A” allele and two spores inherit the “B” allele. Since this information is not explicitly used in calculating the delta scores (which reflect the dependencies between the spore genotypes, but not the exact form of the dependencies), it can be used as a subsequent validation test. For a candidate tetrad to be validated and labeled as a real tetrad, we require the fraction of markers with 2:2 segregation to be close to 1 (above 0.9 by default). For a partial, 3-spore tetrad the process is similar, but the method uses 2:1 segregation and a cutoff of 0.95. We refer to the process of identifying candidate tetrads, performing validation, labeling real tetrads and removing them from the pool of spores used for further consideration as *tetrad verification*.

### Software implementation

Our software implementation combines the previously described methods as follows (Figure 5). First, spore genotypes are preprocessed to remove duplicate spores (identified using the edit-distance between allele vectors) and spores with a large number of missing values. Next, the remaining spores are clustered based on their centromere-flanking markers. Then, the software searches for real tetrads within each cluster based on the delta score of each spore group. Afterwards, all remaining spores that have not been assigned to a tetrad are analyzed to identify any remaining tetrads and then any *partial tetrads*, i.e. groups of three sister spores (with one spore missing from the dataset) and groups of two sister spores (with two spores missing from the dataset). Finally, spores are assigned tetrad labels and output to the user. A detailed flow chart of the search is presented in **Appendix D** (Figure S4).

**Figure 5.**
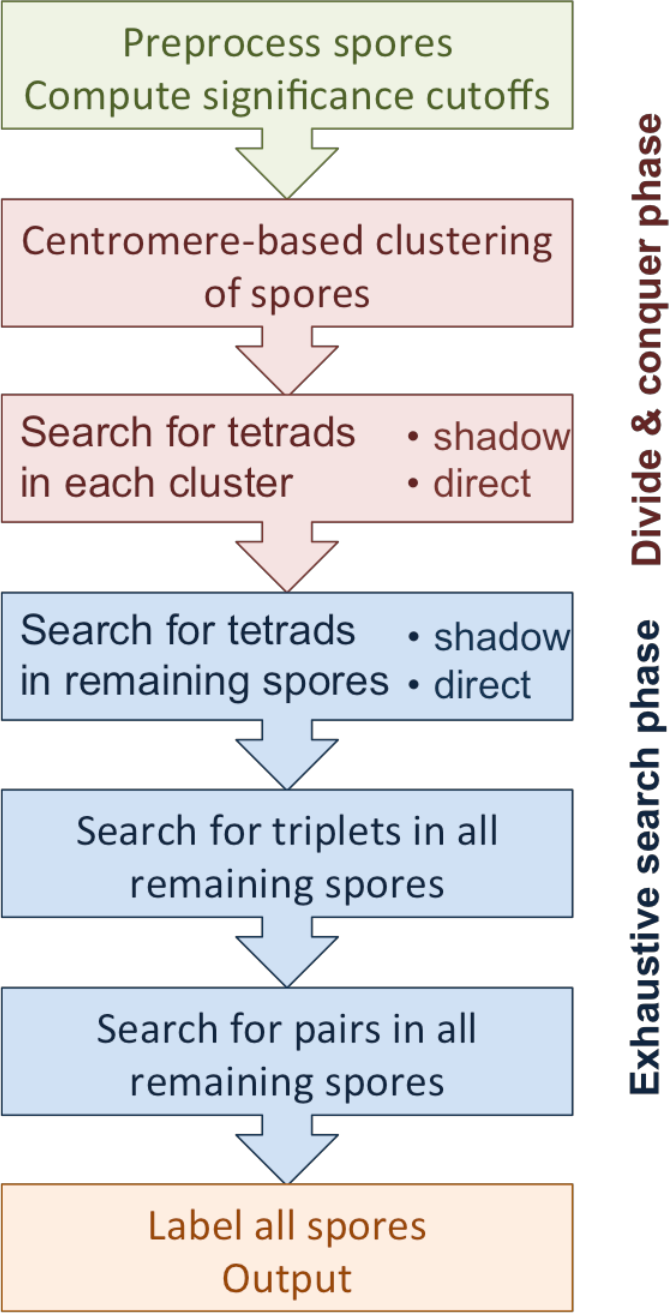
A general overview of the flow of the software for tetrad detection.

The default method for identifying true tetrads within each centromere-cluster is to carry out a *direct* search of all possible combinations of 4 spores using delta. Any set of 4-spore combinations passing the delta score threshold then undergoes the tetrad verification process and, if it passes, is labeled a tetrad and removed from further analysis. The search then continues among the remaining spores. However, for large clusters, exhaustively combing the 4-spore search space in this way is computationally expensive. Therefore, for clusters containing over five spores, tetrads are instead identified in an *indirect*, two-step process. First, because the 3-spore search space is much smaller than the equivalent 4- spore space, the software computes the delta measure on all possible 3-spore combinations and detects those that could be part of true tetrads (hereafter *triplets*). Next, the remaining single spores in the cluster are added to each triplet, and the delta scores are recalculated. Any 4-spore groups that pass the delta score threshold and tetrad verification process are then labeled a full tetrad and removed from further analysis.

The indirect search method described above is an example of a *shadow* search (Sakhanenko and Galas, 2015). The approach leverages the fact that a functional dependency of *N*- variables usually has a detectable signal at a lower degree (fewer variables). For tetrad detection with the delta measure, this is in fact the case, since triplets (3-spore groups that are subsets of a real tetrad) have strong delta scores (Figure 4d and Figure S6a). Therefore, full tetrads can be assembled by identifying high scoring 3-spore subsets within each cluster and then identifying which of the remaining spores belong in a tetrad with each 3-spore group. While this method is more computationally efficient, it relies on the ability of the 3-spore delta measure to distinguish groups of spores from the same tetrad from incorrect groupings, and this discrimination is not as accurate as the one that uses delta calculated on 4-spores (Figure 4b vs 4d). Thus, when computationally feasible, the exhaustive 4-spore approach should be used.

After the direct or indirect identification of tetrads within the centromere clusters, the software searches for tetrads in which only three of the four spores have been placed in the same initial cluster (likely due to noise or a small number of markers). This is also done using the shadow approach. The software computes the delta measure on all possible combinations of three spores in the centromere cluster. Then, 3-spore groups that pass the significance filter are combined with spores from the set of unclustered spores. The software computes the delta measure on these new sets of four spores and performs tetrad verification.

Once every cluster of spores from the centromere clusters has been analyzed, the software moves on to an exhaustive search for remaining tetrads by collecting all the unlabeled spores into a single cluster and applying a combination of shadow and direct searches. If at any point in the search a candidate tetrad fails the verification step based on 2:2 marker segregation, its spores are put back into the search space. This approach is then repeated for triplets, consisting of three sister spores. Finally, any remaining pairs of two sister spores are identified using mutual information, leaving only the unclassifiable single spores.

The software takes as an input a tab delimited text file containing the genotype of each spore. The parameters of the software are defined in a configuration file and preset to the most frequently used values by default. A user can adjust these values in the configuration file to get a better performance for specific situations. Some of the parameters of interest are

- CEN_CALLING controls whether to estimate the recombination frequency from the data or skip it and use the *S. cerevisiae* default specified by COS_PER_MEGA. Estimating recombination frequency on large datasets could be slow, so one might want to skip it, use a published value, or estimate it once before varying other parameters.
- CLUSTERING controls whether we employ the centromere-based clustering first, or go straight to the exhaustive search.
- SIMILARITY_COEFFICIENT is a threshold for centromere similarity between two spores, controlling when the software calls two spores similar. Using the similarity coefficient, we are able to considerably reduce the search space speeding up the processing of large data sets. Smaller centromere similarity coefficient cutoffs result in larger numbers of smaller clusters. Increasing this cutoff will merge the clusters together until they become one cluster – the entire data set. The user should set the cutoff such that there are many medium size (under 300 spores) clusters.
- D4_PVALUE and D3_PVALUE specify the cutoffs for the p-value of the Delta4 and Delta3 scores. To avoid false positives, the user can increase the stringency of the search by lowering the p-value of the Delta4 and Delta3 scores. The user can also adjust the segregation cutoffs (SEG_CUTOFF_D4 and SEG_CUTOFF_D3) to filter out the false positives.

The full list of the parameters with their descriptions is given in the *readme* file of the software. The software also comes with the set of examples described in this paper and the corresponding configuration files to process these examples. The information-theory-based dependency detection is implemented in *C*, whereas the rest of the software is implemented in *Python*. The software is available for download here: *URL*

### Testing the software on simulated and real datasets

We first applied the software to simulated datasets. We generated three sets of error-free *S. cerevisiae* tetrad genotypes (see **Appendix F** for more details), including recombination events and recording spore relationships: *Small* (100 tetrads and 1,000 markers), *Medium* (1,000 tetrads and 10,000 markers), and *Large* (1,000 tetrads and 100,000 markers). For each of these datasets, we created multiple test cases by reducing the number of markers and adding various amounts of missing values and levels of noise (Table 1).

**Table 1.** The tetrad detection software applied to various simulated test sets. Three simulated datasets are shown (indicated in the column Set): *Small* (in blue, 100 tetrads, 1000 markers), *Medium* (in yellow, 1000 tetrads, 10000 markers), and *Large* (in red, 1000 tetrads, 100000 markers). The first four columns of the table show the various parameters of the simulated data considered: N_t_ shows the number of tetrads in the set, N_m_ shows the number of randomly selected markers as well as the percentage of the original set of markers, and columns Noise and Missing show the amount of noise and missing values (in percent) added to the data. The software was run on these test sets with different parameters. The table shows only the parameter settings that produced the optimal result. For the results of the software with other settings see Table S1 in **Appendix H**. Column “Cent calling” shows whether the software estimates the recombination frequency empirically or uses its default value derived from published data (Mancerra et al. 2008). The column “Sim coeff.” shows the similarity threshold being used. The columns 2:2 cutoff and 2:1 cutoff show the cutoff values for 2:2 and 2:1 segregation score for the software to call two spores sisters. The last two columns show the performance of the software, the total runtime (Intel Core i7-7820X CPU @ 3.60GHz and 64 GB RAM) and the percent of total tetrads detected by the software. Note that in all the cases when the software detected 100% of tetrads, there were no false positives. In the case when 99.8% of tetrads were detected, there was 1 false positive tetrad and 1 false positive triplet (with 1 spore unassigned).

| Set | Test set parameters |  |  |  | Software parameters |  |  |  | Performance |  |
| --- | --- | --- | --- | --- | --- | --- | --- | --- | --- | --- |
|  | N<sub>t</sub> | N<sub>m</sub> | Noise % | Missing % | Cent. calling | Sim coeff | 2:2 cutoff | 2:1 cutoff | Run time | Detected tetrads % |
| **Small** | 100 | 500 (50%) | 0 | 0 | yes | 0.21 | 0.9 | 0.95 | 6.0 | 100% |
|  |  |  | 0 | 5 | yes | 0.19 | 0.9 | 0.95 | 5.7 | 100% |
|  |  |  | 0 | 10 | yes | 0.2 | 0.9 | 0.95 | 6.2 | 100% |
|  |  |  | 5 | 0 | yes | 0.2 | 0.75 | 0.88 | 6.7 | 100% |
|  |  |  | 10 | 0 | yes | 0.24 | 0.6 | 0.83 | 8.1 | 100% |
|  |  | 984 (100%) | 0 | 0 | yes | 0.13 | 0.9 | 0.95 | 12.9 | 100% |
|  |  |  | 0 | 5 | yes | 0.14 | 0.9 | 0.95 | 12.3 | 100% |
|  |  |  | 0 | 10 | yes | 0.13 | 0.9 | 0.95 | 12.1 | 100% |
|  |  |  | 5 | 0 | yes | 0.2 | 0.75 | 0.88 | 14.3 | 100% |
|  |  |  | 10 | 0 | yes | 0.22 | 0.6 | 0.83 | 16.0 | 100% |
| **Medium** | 1000 | 1000 (10%) | 0 | 0 | no | 0.12 | 0.9 | 0.95 | 02:10.2 | 100% |
|  |  |  | 0 | 5 | no | 0.11 | 0.9 | 0.95 | 02:37.4 | 100% |
|  |  |  | 0 | 10 | no | 0.12 | 0.9 | 0.95 | 02:27.2 | 100% |
|  |  |  | 5 | 0 | no | 0.14 | 0.75 | 0.88 | 27:15.2 | 100% |
|  |  |  | 10 | 0 | no | 0.16 | 0.6 | 0.83 | 14:14:47.7 | 99.80% |
|  |  | 9984 (100%) | 0 | 0 | no | 0.1 | 0.9 | 0.95 | 03:38.2 | 100% |
|  |  |  | 0 | 5 | no | 0.1 | 0.9 | 0.95 | 04:09.1 | 100% |
|  |  |  | 0 | 10 | no | 0.09 | 0.9 | 0.95 | 03:32.5 | 100% |
|  |  |  | 5 | 0 | no | 0.12 | 0.75 | 0.88 | 17:35.0 | 100% |
|  |  |  | 10 | 0 | no | 0.17 | 0.6 | 0.83 | 26:43:15.6 | 100% |
| **Large** | 1000 | 10000 (10%) | 0 | 0 | no | 0.07 | 0.9 | 0.95 | 03:28.3 | 100% |
|  |  |  | 0 | 5 | no | 0.12 | 0.9 | 0.95 | 03:40.0 | 100% |
|  |  |  | 0 | 10 | no | 0.09 | 0.9 | 0.95 | 03:35.7 | 100% |
|  |  |  | 5 | 0 | no | 0.12 | 0.75 | 0.88 | 18:46.4 | 100% |
|  |  |  | 10 | 0 | no | 0.16 | 0.6 | 0.83 | 31:13:42.7 | 100% |
|  |  | 50000 (50%) | 0 | 0 | no | 0.07 | 0.9 | 0.95 | 10:45.2 | 100% |
|  |  |  | 0 | 5 | no | 0.05 | 0.9 | 0.95 | 08:33.1 | 100% |
|  |  |  | 0 | 10 | no | 0.07 | 0.9 | 0.95 | 10:40.6 | 100% |
|  |  |  | 5 | 0 | no | 0.12 | 0.75 | 0.88 | 36:59.5 | 100% |
|  |  |  | 10 | 0 | no | 0.14 | 0.6 | 0.83 | 93:53:41.7 | 100% |

We used this simulated data to perform a thorough evaluation of the individual components of the software in various situations. Specifically, we analyzed the performance of allele calling at centromeres, spore clustering based on similarity coefficients, and tetrad verification using segregation scores. **Appendix C** shows the details of the evaluation on the simulated dataset *Small*.

In general, the total number of markers in the data affected the estimation of the allele calls at centromeres and consequently the initial clustering of the spores. Low numbers of markers caused problems in clustering through several mechanisms. These included poor precision in estimating global recombination parameters and difficulties in accurately estimating the allele at the centromere due to a lack of flanking markers or because the flanking markers were too far apart.

The proportion of missing data, on the other hand, did not have a strong effect on centromere allele call predictions or spore clustering, and any effect was fully overcome by using more markers. The noise in the data (genotyping errors), however, had a notable impact on the performance of the software. High levels of noise strongly affected the estimation of the alleles at the centromeres and spore clustering (**Appendix C**). This problem can, however, also be overcome by increasing the number of markers used. Furthermore, the noise affected the strength of the true tetrad signal relative to the background, making it harder to distinguish the true tetrads statistically from the other 4- spore sets. We note that this was the case for both the delta measure as well as the 2:2 segregation measure. The quantitation of this effect is described in **Appendix C**.

After testing the individual components of our software in this way, we then tested our software on a handful of simulated test sets derived from the sets *Small*, *Medium*, and *Large* and compared the final tetrad assignments to ground truth from the simulations. All the tests were performed on a desktop with Intel Core i7-7820X CPU @ 3.60GHz (8 cores, 16 threads) and 64 GB RAM. Table 1 summarizes these tests and shows the corresponding optimal parameter settings of the software and the resulting performance. A comprehensive list of the tests performed is presented in Table S1 of **Appendix H**.

Table 1 shows that the software is able to achieve 100% accuracy with no false positives in almost all the tests: only in tests on the *Medium* set with only 10% of the original marker set and with 10% noise was the accuracy lower, albeit still 99.8%. In these tests some 4- spore groups scored as high as the real tetrads due to noise in the relatively small number of markers, resulting in two missed tetrads, one false positive tetrad assignment, and one false positive triplet assignment.

The runtime of all the tests without noise was very reasonable: *Small* tests took under 12 seconds, *Medium* tests took under 4 minutes, and *Large* tests took under 11 minutes. Although with the addition of noise, the runtime of *Small* tests did not change much due to the small size, it changed drastically for *Medium* and *Large* tests. At 5% noise, *Medium* and *Large* tests took under 28 minutes and 37 minutes correspondingly. Increasing noise to 10% made the runtime go up considerably: *Medium* tests took 14-27 hours (depending on the size) and *Large* tests took 31-94 hours. In general, including more markers in the test sets increases the time it takes to calculate each delta score. Without noise, the clustering of spores is very effective, allowing for the search space to be divided into multiple small clusters fully containing tetrads, thus keeping the number of delta calculations low and resulting in a short runtime. With noise however, the initial clustering does not work as well, which inevitably increases the total number of delta calculations necessary and thus increases the runtime.

To achieve the best performance, we varied two software parameters, the similarity coefficient threshold and the segregation cutoffs. In test cases where the number of markers was large, the default similarity coefficient threshold resulted in a small number of very large clusters, causing the software to take a considerable amount of time to complete a run (**Appendix C**). Lowering this threshold increased the number of clusters while reducing their size, which divided the search space more efficiently and resulted in faster processing. Users of this software should consider these options, since the level of noise in experimental datasets is usually not known.

When no noise is present, clustering allows the software to subdivide the spores into sets of same-tetrad spores, identifying all the tetrads at the divide-and-conquer phase (see Figure 5). In this situation, keeping the clusters small results in smaller processing time. For the test cases with noise however the clustering does not work as well and, as a result, only some tetrads are identified at the divide-and-conquer phase leading to more time spent in the exhaustive search phase (see Figure 5). In this case making the clusters larger allows the software to extract more tetrads at the divide-and-conquer phase, thus reducing the amount of computation at the exhaustive search phase and the overall processing time. Therefore, for the cases with noise (in particular when the noise level is 10%) the similarity threshold had to be increased relative to the same sized dataset without noise resulting in larger clusters. Furthermore, in the presence of noise the default values of the segregation cutoffs are too stringent, resulting in poor accuracy, since many true tetrads were dismissed. See **Appendix C** for more details about the effects of the similarity coefficient and the segregation cutoffs.

We then applied the software to real biological data from two published datasets. Dataset D1 consists of 412 spores from manually dissected tetrads and the tetrad relationship between spores was recorded by the experimenter (Sirr, et al. 2017). Dataset D2 consists of 3,200 spores that were randomly arrayed on an agar plate, but that contained a tetrad-specific plasmid barcode (Ludlow et al. 2013). In both cases, the genotype markers were generated by RAD-seq as described in the original publications (Sirr, et al. 2017; Ludlow et al. 2013). Using these datasets allowed us to compare the effectiveness of our method to the experimentally derived tetrad assignments, which we treated as “ground truth” (Table 2).

**Table 2.**
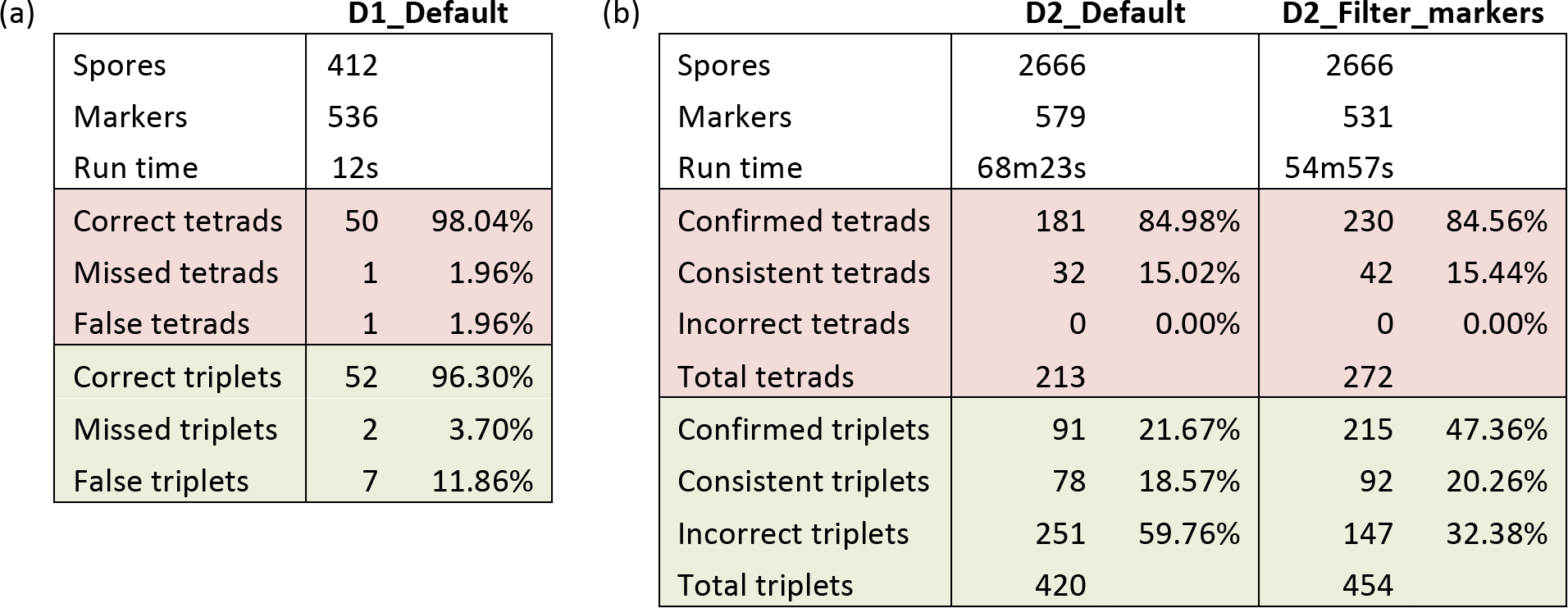
The tetrad detection software applied to real biological datasets. (a) Evaluation of the software on dataset D1 using the default parameter setting. Since all the spores in D1 are already labeled according to their tetrad assignment (from manual dissection), we were able to count the number of tetrads identified by the software that were either in agreement with the data labels (correct tetrads), or not (false tetrads), as well as the number of tetrads that the software was not able to identify (missed tetrads). The triplets were counted in the same way. (b) Evaluation of the software on dataset D2 using either the default parameter setting or filtering markers with too many missing values (corresponding to the second and third columns). Although there is no known tetrad assignment of the spores in D2, the spores were barcoded, with spores derived from the same tetrad sharing a barcode. As a result we counted the number of tetrads identified by the software comprised of same-barcode spores (confirmed tetrads). We also counted consistent tetrads comprised of either same-barcode spores or spores with no barcode and incorrect tetrads whose spores have different barcodes. The sum of confirmed, consistent and incorrect tetrads is the total number of tetrads identified by the software. The triplets were counted in the same way.

For dataset D1, using the default parameters the software achieved over 98% full-tetrad accuracy and over 96% accuracy of identifying triplets (tetrads missing 1 spore). For dataset D2, after filtering out the duplicate spores, our software with default parameters showed 100% agreement between identified tetrads and the barcodes (sum of number of confirmed and consistent tetrads from Table 2). However, almost 60% of triplets identified by the software in default mode were in disagreement with the barcodes. Filtering to remove markers with large numbers of missing values improved the 3-spore performance on dataset D2 by almost 30%. Note also that filtering markers allowed identifying considerably more tetrads and triplets.

### Data availability statement

Supplemental files available at FigShare. File tetrad_detection_software.rar contains the code of the tetrad assembly software, as well as code used to generate the simulated data, the information files and the test data. File simulated_data.rar contains all the simulated data used in the paper to test our software.

## Discussion

We have described a computational method, based on our previously developed information theory dependency analysis, that reconstitutes tetrads using only information from genome sequencing of the meiotic progeny. This method avoids the need for tetrad-specific barcoding and or genetic modification of any kind. Instead, our method uses information associated with specific features of tetrad genome sequences, features that result from the mechanisms of meiosis. The software reported here is both simple to use and highly effective in reconstituting tetrads. This approach can significantly increase the power of tetrad analysis in several ways, most notably by vastly increasing the numbers of tetrads that can be analyzed with minimal effort.

Our software integrates a heuristic method for clustering, using centromere proximal markers, with an information theory-based method for signal detection. On both simulated and real data, the software achieves a remarkably high success rate, even in the presence of lost spores and sources of experimental noise. The applications for this software include, the analysis of meiotic recombination and the study of gene conversions or other non-reciprocal genetic events in which a subset of markers deviate from the expected 2:2 segregation pattern.

The software can also be applied to any organism in which it is possible to isolate the four products of meiosis. Our software is based on an information theory method, which does not make any assumptions about the data and its underlying structure. Although the genome structure of *S. pombe* is significantly different from that of *S. cerevisiae*, the analysis method described here will work equally well on this organism. Because the software only uses the centromere allele patterns of the organism to reduce the search space and thus decrease run times, it can be effectively applied to organisms with smaller numbers of chromosomes. For example, in *S. pombe*, which has only three chromosomes, the method would still produce several, albeit large, clusters of spores and thus productively divides the search space.

## Appendix A.

In the analysis of complex biological systems we need measures that can detect synergistic, multiple variable dependencies. Mutual information is a well-known measure that quantifies the amount of dependency between two variables:

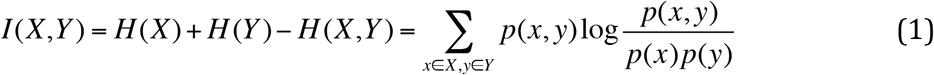

Interaction information has been proposed (McGill, 1954) as a multivariable generalization of mutual information. This measure has a number of advantages and drawbacks (Bell, 2003; Jakulin and Bratko, 2004; Sakhanenko and Galas, 2011) but can be used to devise powerful measures of dependency for any number of variables. The interaction information for three variables, for example, quantifies the difference between the two-variable interaction information (mutual information), with and without knowledge of the third variable:

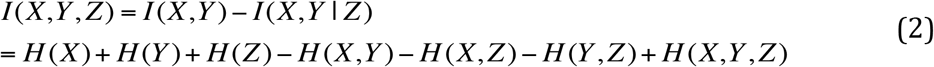

Here*I*(*X*,*Y* | *Z*) is conditional mutual information, *H* (*X*) is entropy of variable *X* and *H* (*X*,*Y*, *Z*) is a joint entropy of the three variables. Note that the conditional mutual information is actually a difference between interaction informations for two and three variables – a *differential interaction information*. A general form of interaction information for the set of ν_*n*_ variables, in terms of marginal entropies can be written as:

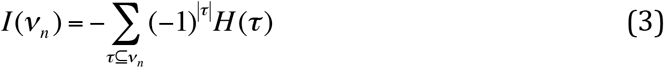

In this paper we use a symmetric product of *differential interaction information*, which we call “delta” (Galas et al., 2014; Sakhanenko and Galas, 2015; Galas and Sakhanenko, 2016). Differential interaction information quantifies the change in interaction information that occurs when we add another variable to a set of variables, so for three variables it is defined as:

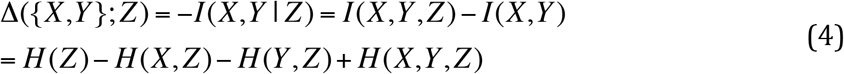

If ν_*n*_ = {*X*_1_, *X*_2_ … *X*_*n*_} and ν_*i*_ = ν_*n*_-{*X*_*n*_} then the differential interaction information can be defined in general as

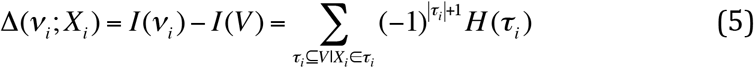

Note that, unlike interaction information, differential interaction information is not symmetric, since *X*_*i*_ in equation 5 is a special variable. In order to create a symmetric measure, we take the product of differential interaction information with all possible choices of the target variable:

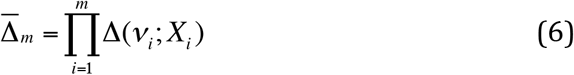

We refer to 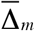 as the *delta* measure, for *m* variables. Although this is a general, multi-variable measure, in this paper we focus on delta computed only on 3- and 4-variable sets. We use 3 and 4-variable *delta*, as well as the pair-wise measure, mutual information, to scan the data from large sets of yeast spores and detect and assemble spore tetrads and their components.

## Appendix B

In order to be able to cluster the spores, we need to define a measure of similarity of two spores based on allelic information at the centromeres. We first start with a deterministic case, when the alleles near the centromeres are known.

Consider two vectors X and Y, whose elements are either 1 or 2, representing allele calls at the centromere for the two corresponding spores. We can then define a vector 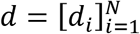 such that *d*_*i*_ = 1 if *X*_*i*_ = *Y*_*i*_, and 0 otherwise, where *N* is the number of centromeres. Note that if we define a vector *b* = [*b*_*i*_] such that *b*_*i*_ = 0 if *X*_*i*_ = *Y*_*i*_, and 1 otherwise, then *b* = 1 − *d*.

Consequently, we define S, the coefficient of similarity of X and Y, as follows

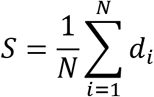

The coefficient of anti-similarity *U* is defined similarly to *S*, but using vector *b* instead. Note that by definition *U* = 1 − *S*.

Given the allele calls at the centromeres, we can compute S for all possible pairs of spores and then cluster the spores based on their similarity at the centromeres. In most biological examples, however, the allele values at the centromeres are unknown. We can however estimate what the allele is at each centromere based on the recombination frequency in a given data set and the allele calls at the markers flanking the centromeres (see **Appendix G**), and consequently we can estimate the coefficient of similarity between spores.

Given the probabilities *P*(*X*_*i*_ = *a*) and *P*(*Y*_*b*_ = *b*) that spores *X* and *Y* have alleles *a* and *b* at the centromere *i*, we can define the probability that *X* and *Y* are identical at *i*th position as *p*_*i*_:

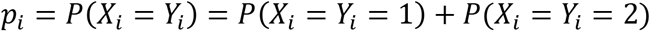

If *X*_*i*_ and *Y*_*i*_ are independent, then the definition above can be simplified as

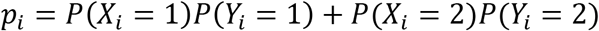

Since we do not know the values of *X* and *Y*, we cannot compute the similarity coefficient directly, thus we need to estimate it. An estimated coefficient of similarity 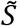 is computed by taking the expectation of *S*:

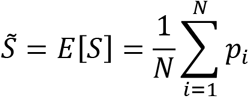

Note that an estimated coefficient of anti-similarity is 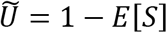.

As an example, we apply this similarity coefficient to the simulated data set of 400 spores and 500 markers with no noise or missing values. For each pair of spores we compute the similarity coefficient based on the centromere allele estimates. Figure S1 shows the distributions of coefficients for the spores from the same tetrad and for the spores from different tetrads.

**Figure S1.**
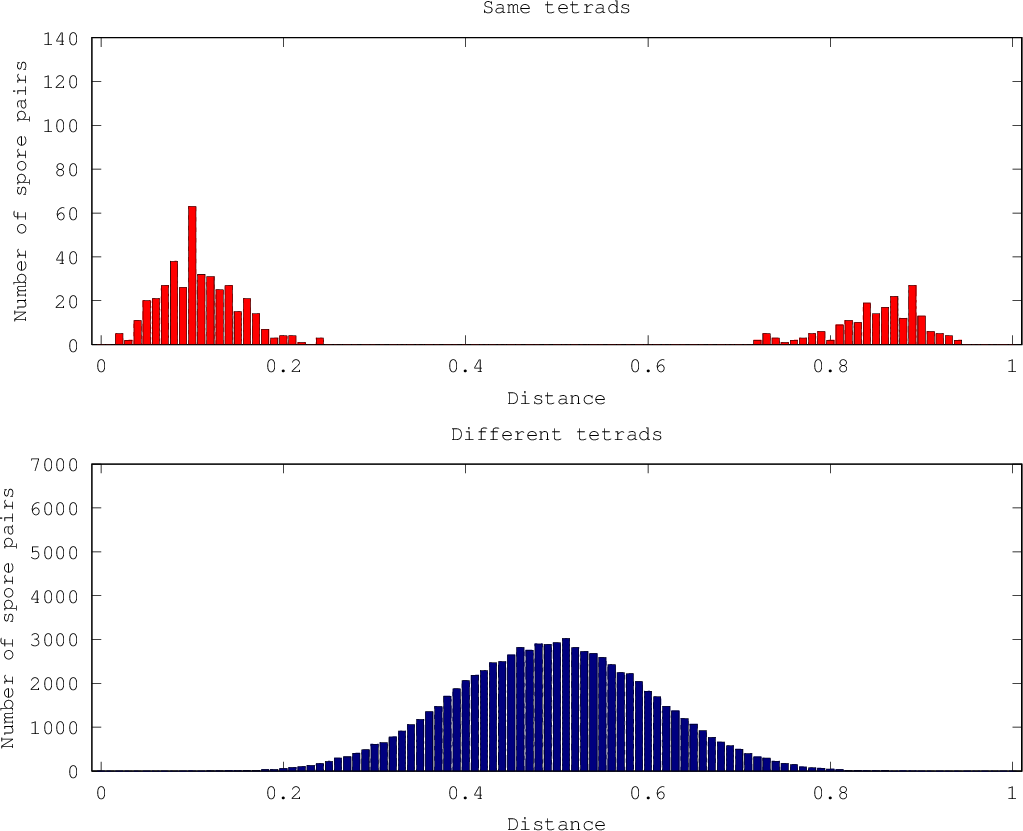
Distribution of estimated similarity coefficients for spore pairs from the same tetrad (in red) and from different tetrads (in blue) computed on the simulated data set (500 markers, 400 spores, 0% noise, 0% missing).

This figure shows the expected pattern that, for centromeric alleles, each tetrad consists of two pairs of identical spores that are reflections of one another. The reflected spores in each tetrad form a distribution near 0, whereas the completely identical spores form a distribution near 1. On the other hand, the spores from different tetrads form a normal distribution centered around 0.5.

Form figure S1 it is clear that we can use the similarity coefficient to cluster the spores from the same tetrads. The distributions from figure S1 can be transformed by “folding” it in on itself as follows, *S*’ = 0.5 − |0.5 − *S*|, such that *S’* is close to 0 when spores are from the same tetrads and around 0.5 when the spores are from different tetrads. The scores in between correspond to situations when the allele call estimates at the centromere are not enough to identify the spores from the same tetrad due to noise and missing data. We need to find a threshold *T* that simultaneously maximizes the number of same tetrad spores with *S*’ < *T* as well as the number of spores from different tetrads with *S*’ > *T*. We determined that *T* = 0.*2* is the optimal threshold to correctly estimate the centromere allele calls when the number of markers is sufficient and the noise and missing data is minimal.

## Appendix C

We now investigate how the noise and missing values affect the components of the software. We generated a number of simulated data sets with varying number of markers and spores, as indicated in the text. We treated the data set as a long vector and added noise by flipping the values of the vector at randomly selected positions. Similarly, we added missing values by erasing the values of the vector at randomly selected positions. We considered different levels of noise and missing values, up to 25% for both.

### Allele calling at centromeres

Using the simulated data we looked at the effects of noise and missing data on the estimation of alleles at the centromeres. As expected, increasing noise makes the centromere allele estimates worse (see Figure S10). On the other hand, the estimates do not seem to be affected by the amount of missing data.

**Figure S10.**
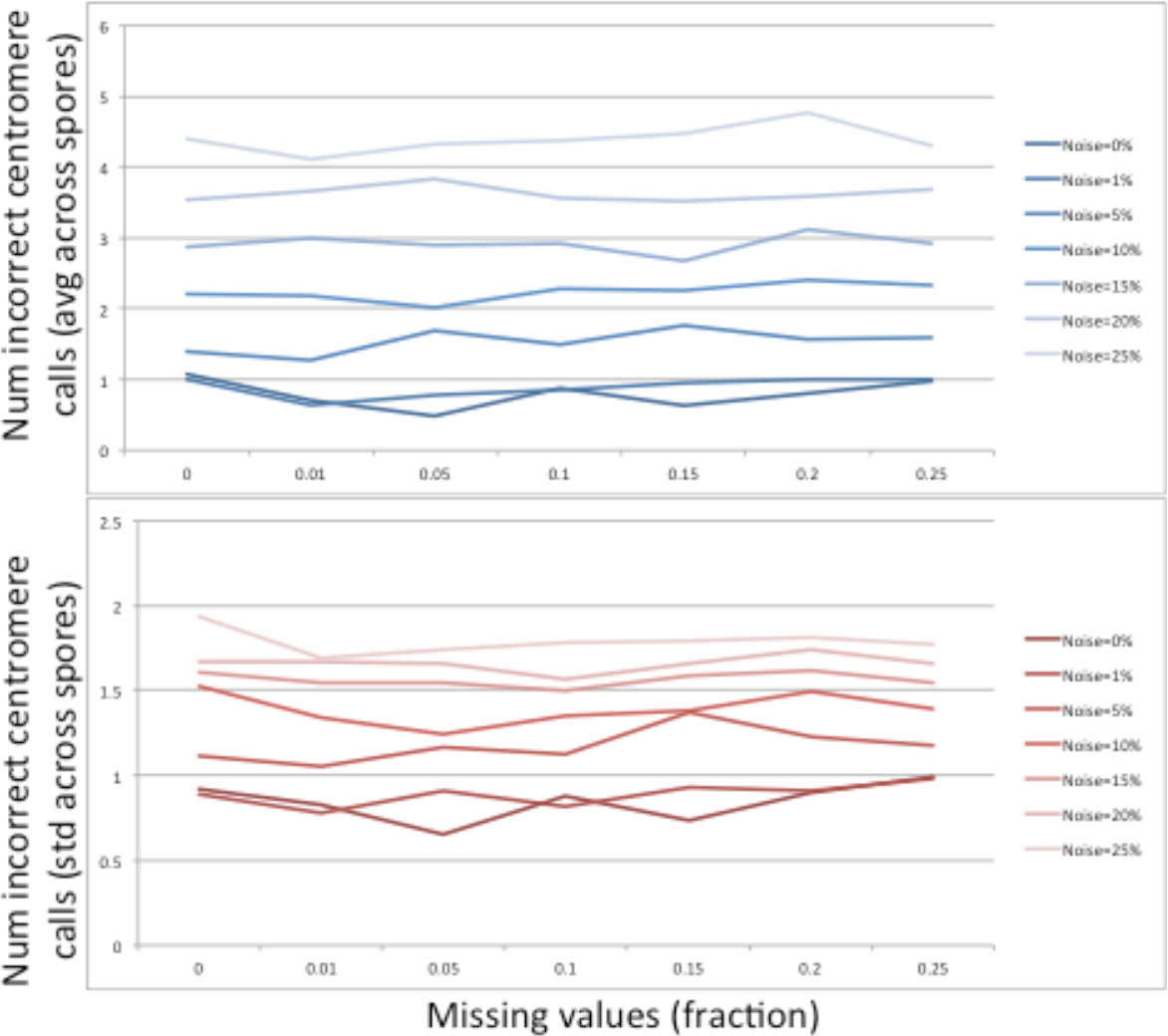
Average (top) and standard deviation (bottom) of the number of incorrect allele calls at the centromeres computed on the simulated data set (500 markers, 400 spores). The x-axis corresponds to the fraction of the missing values in the data set. Each line corresponds to the noise level, ranging from 0% to 25%.

We also observed that a smaller number of markers makes the estimate worse. This is because with fewer markers, some chromosomes either lack flanking markers or their flanking markers are very far apart, making predicting centromere alleles difficult. Figure S11 shows that as the distance between flanking markers increases, the number of incorrect allele calls tends to increase.

**Figure S11.**
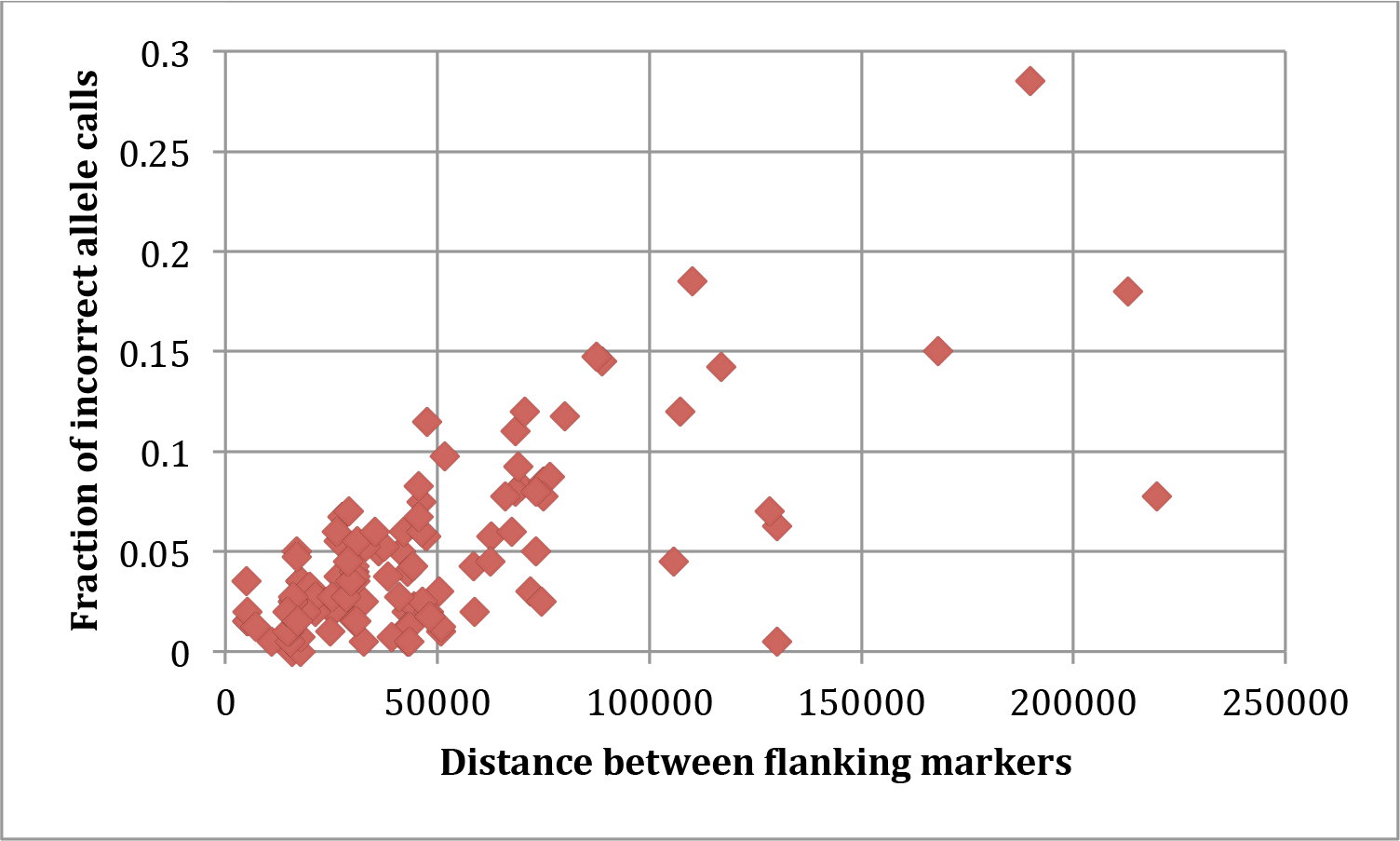
The distance between the flanking markers versus the fraction of incorrect allele calls for each centromere computed on the simulated data set (500 markers, 400 spores, no noise, and the fraction of the missing values ranging from 0% to 25%).

### Similarity coefficients

We now look at the similarity coefficients. Since the centromere allele estimates become less precise when too few markers are selected, it is expected that the centromere similarity coefficients will also not perform well. Figure S12 shows that the range of similarity scores narrows when fewer markers are used and the coefficient distribution of spores from the same tetrads starts to overlap that of spores from different tetrads, making it harder to cluster the spores. Note that, on the one hand, a value around 0.2 is optimal in the case of 500 markers, but on the other hand, it is not even nearly optimal in the case of 100 markers.

**Figure S12.**
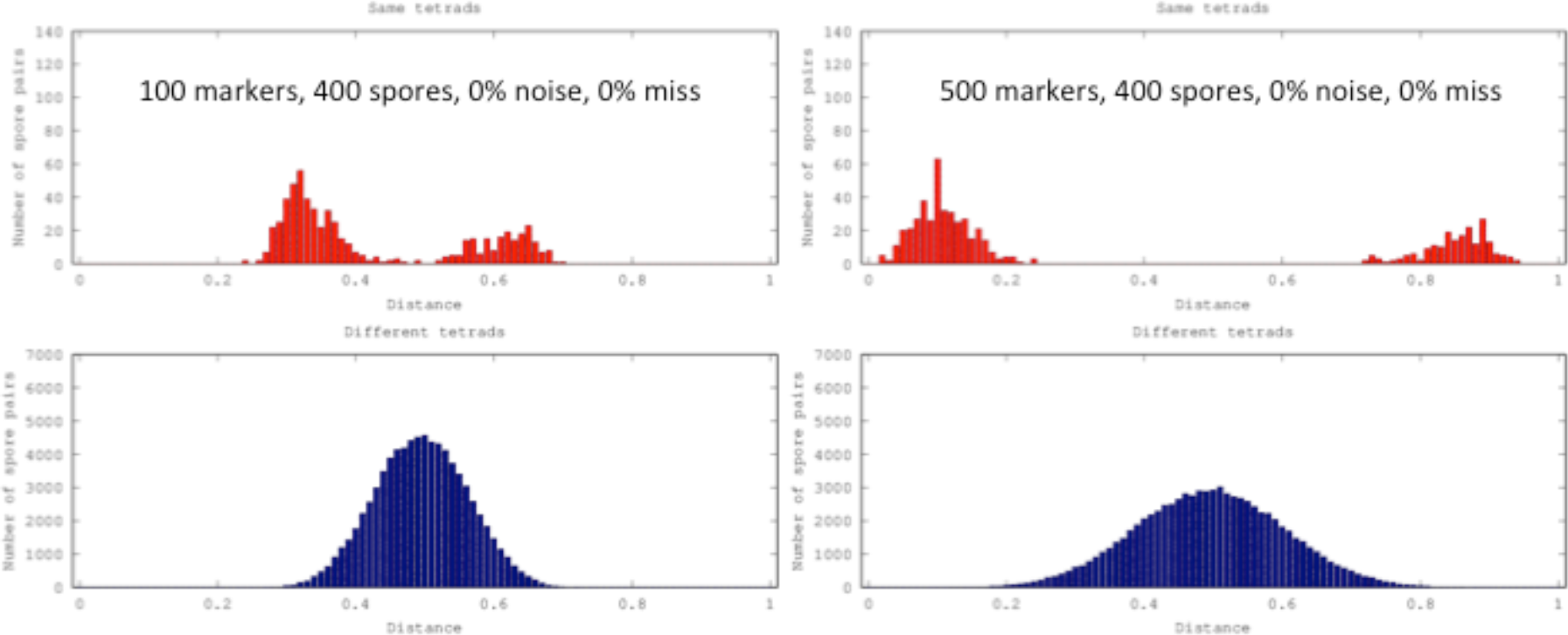
Distributions of centromere similarity coefficients between pairs of spores from the same tetrad (top panels) and from different tetrads (bottom panels). Left panels show the similarity coefficients computed from the simulated data set consisting of 400 spores and 100 randomly selected markers, whereas 500 markers were selected to compute the coefficients in the right panels. The data set had no noise or missing values.

A similar effect is observed when we add noise to the data set. Figure S13 shows that the distribution of similarity coefficients for spores from the same tetrads gradually moves from the extremes towards the middle and becomes indistinguishable from the distribution of spores from different tetrads. Note that the 0.2 threshold is still usable when the noise level is relatively low, and stops working when the noise is over 5%.

**Figure S13.**
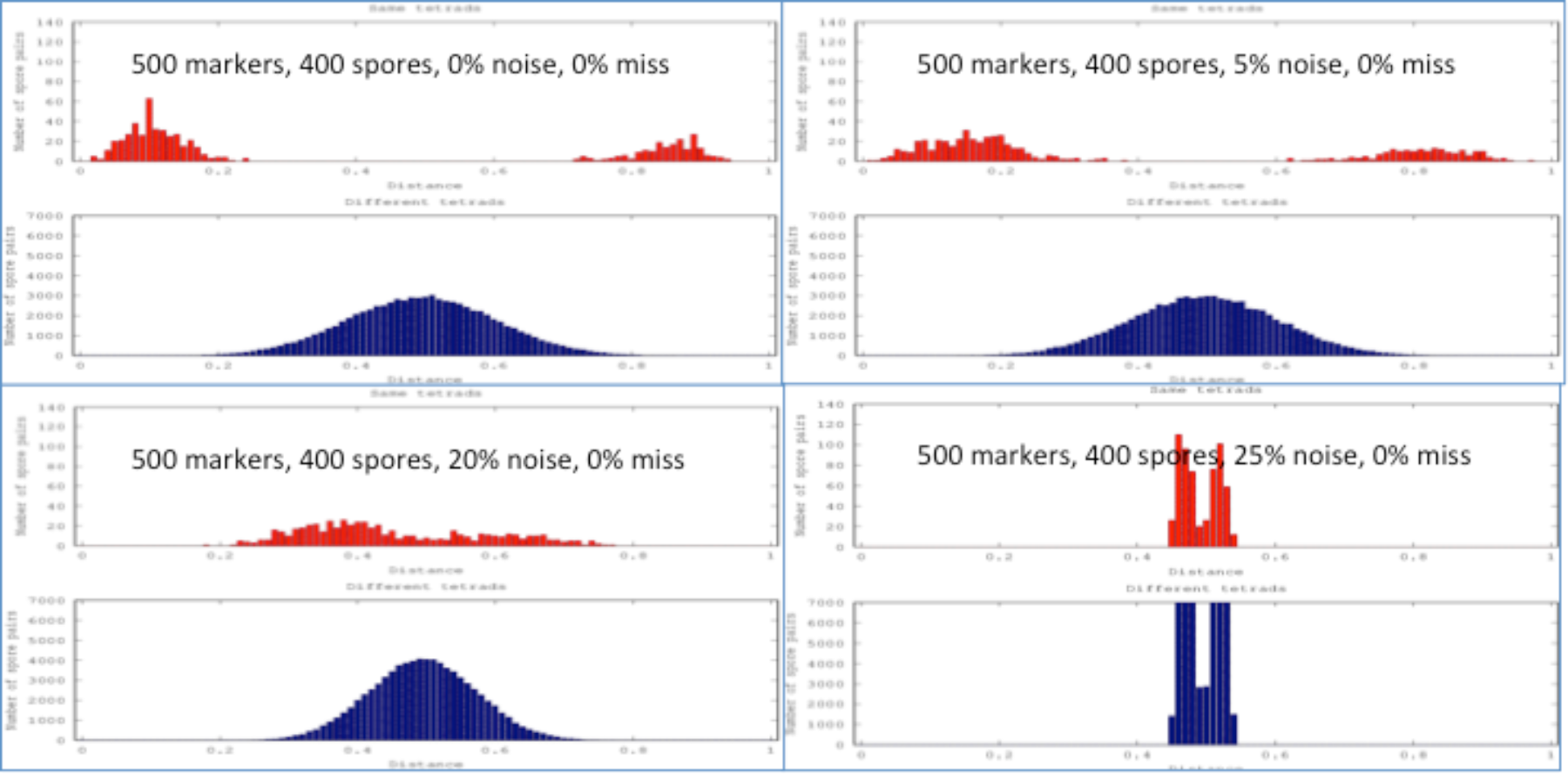
Distributions of centromere similarity coefficients between pairs of spores from the same tetrad (in red) and from different tetrads (in blue). The simulated data of 500 markers and 400 spores was generated at 4 different levels of noise, 0%, 5%, 20%, and 25%. The data sets had no missing values.

On the other hand, as Figure S14 shows, missing data generally does not have a significant effect on the similarity coefficients. Moreover, this is the case even when rather few markers are considered (100 markers).

**Figure S14.**
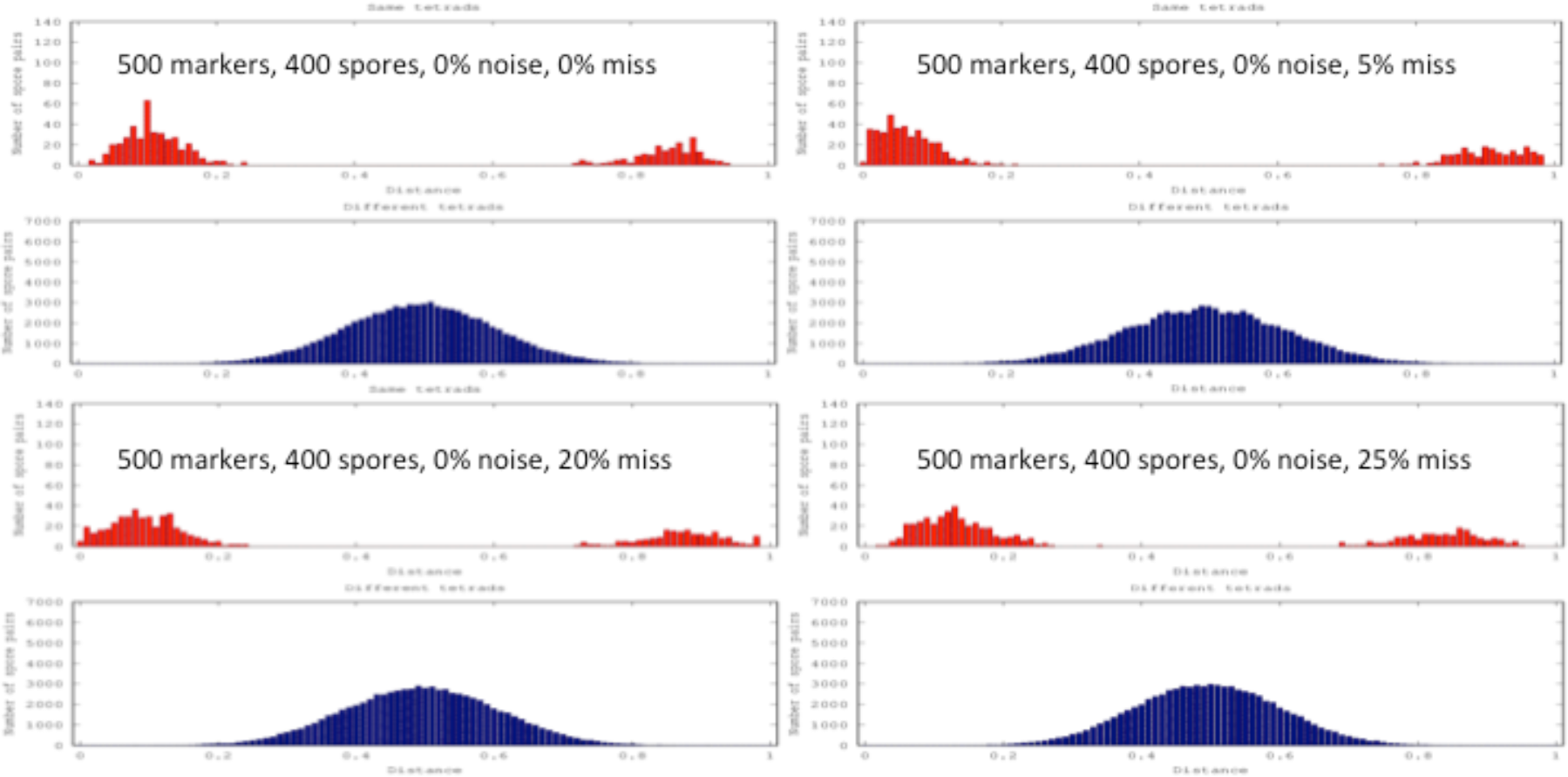
Similar to Figure 13, but this time the simulated data was generated using 4 different levels of missing information, 0%, 5%, 20%, and 25%., and no noise was added.

### Segregation scores

For a 4-spore set, the 2:2 segregation score measures the fraction of genetic positions where two spores have allele “A” and the other two spores have allele “B”. Therefore, for a perfect tetrad the 2:2 segregation score is 1. This changes however when we add noise to the data set. If *p* is the probability of a genetic position having noise (probability of a flipped value), then the 2:2 segregation score of a tetrad is

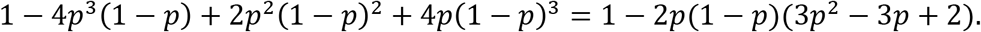

Similarly, for a 3-spore subset of a tetrad, the 2:1 segregation score decreases from 1 to: 1 − *p*^2^(1 − *p*) + *p*(1 − *p*)^2^ = 1 − *p*(*p* − 1). Note that both 2:2 and 2:1 segregation scores are minimal when *p* = 0.5. Table S1 shows the 2:2 and 2:1 segregation scores for true tetrads under different levels of noise.

**Supplementary Table S1.**
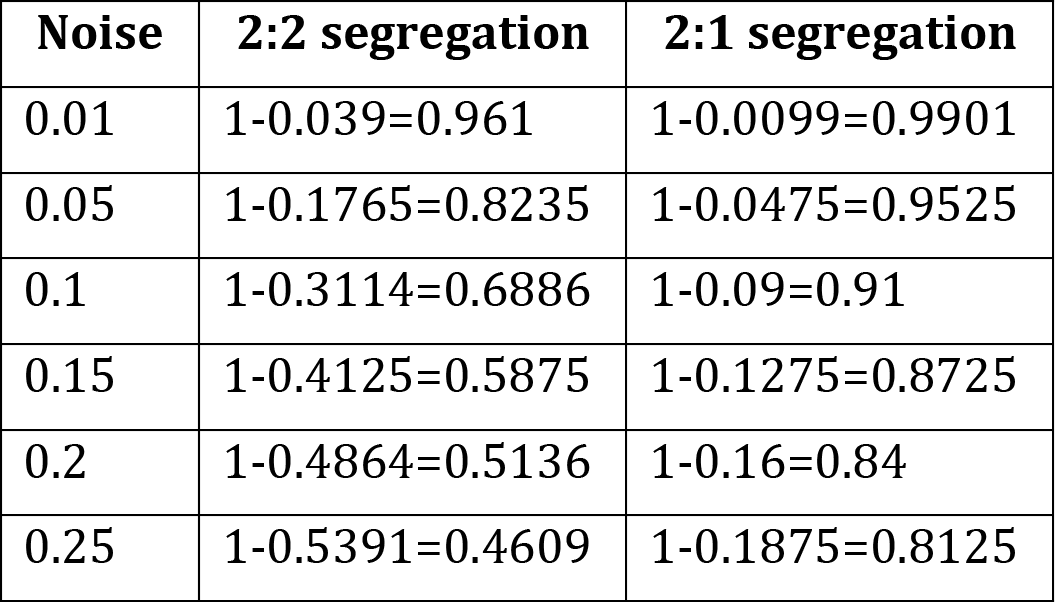
The change of the 2:2 and 2:1 segregation values depending on the noise level.

| Noise | 2:2 segregation | 2:1 segregation |
| --- | --- | --- |
| 0.01 | 1-0.039=0.961 | 1-0.0099=0.9901 |
| 0.05 | 1-0.1765=0.8235 | 1-0.0475=0.9525 |
| 0.1 | 1-0.3114=0.6886 | 1-0.09=0.91 |
| 0.15 | 1-0.4125=0.5875 | 1-0.1275=0.8725 |
| 0.2 | 1-0.4864=0.5136 | 1-0.16=0.84 |
| 0.25 | 1-0.5391=0.4609 | 1-0.1875=0.8125 |

## Appendix D

The tetrad software consists of four main components: the preprocessing steps, the heuristic search, the direct search, and the post-processing steps (see Figure S4).

**Figure S4.**
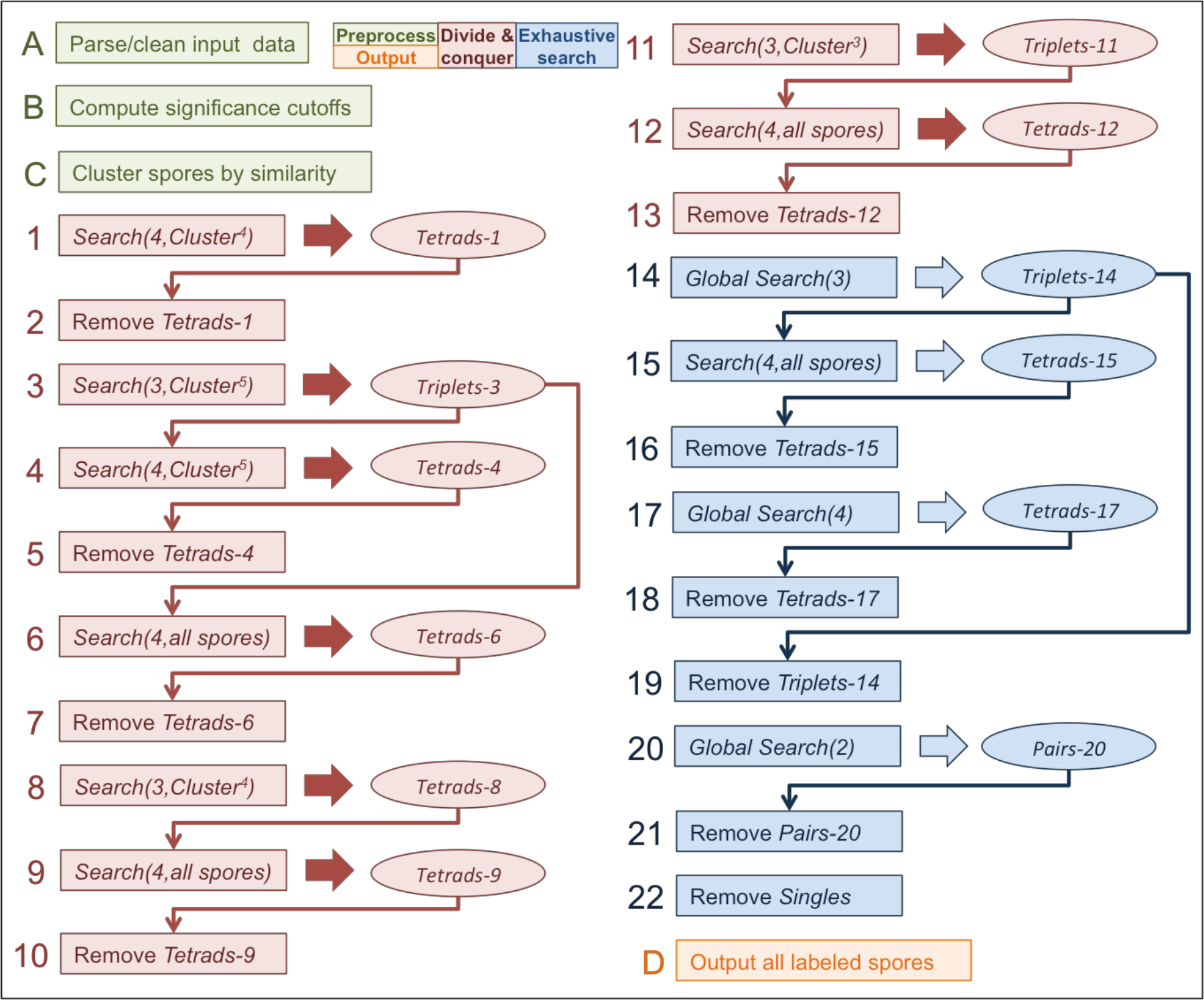
Detailed view of the software for tetrad detection. The flow is shown sequentially top to bottom, left to right, with the steps labeled with a letter/number. Four different components of the software are identified with different colors (see the legend on the top of the figure): the preprocessing is in green, the heuristic search is in red, the direct search is in blue, and the post-processing is in yellow. The general functions are shown in the boxes with their resulting sets of tuples shown in ovals. The thin arrows show where the tuple sets are being used as inputs to functions. Most of the general functions shown are self-explanatory. Function *Search(N,Set)* searches for *N*-tuples of spores from *Set*. A thin arrow entering *Search(4,Set)* function indicates that this is a shadow search that traverses all possible quads of spores consisting of a triplet identified by the arrow and a spore from *Set*. Function *Global Search(N)* corresponds to a search for *N*-tuples among all the remaining spores. Set *Cluster*^*N*^ in the argument of a search function traverses all clusters of spores with size N=3, 4, or 5+.

The preprocessing component (steps A-C in green in Figure S4) reads the data in and cleans it by removing spores with too many missing data points, while flagging any duplicate spores. At the next step, the software pre-computes the significance thresholds – the delta scores that correspond to a user-defined level of significance (*p-value*=0.05 by default set by the software parameters D4_PVALUE, D3_PVALUE, D2_PVALUE) for quads, triplets, and pairs. These thresholds correspond to the minimal delta scores required for a tuple to be considered a real tetrad or part of a real tetrad. To compute the thresholds, the software generates a set of *N* random tuples set by the parameter RANDOM_SAMPLE_SIZE (equal 10000 by default), calculates delta scores, and finds the score corresponding to the preset p-value. Finally, the preprocessing component attempts to cluster the spores based on their centromere similarity (the user may choose to switch the clustering off by setting the parameter CLUSTERING to 0). To cluster the spores, the software first estimates the recombination frequency as a function of physical distance (see **Appendix G**). On large datasets this step can take too long, therefore the user can skip it (by setting CEN_CALLING to 0) and use the default value of the conversion factor (COS_PER_MEGA=3.7) derived from (Mancera et al, 2008). Given the recombination frequency, the physical position of centromere-flanking markers and the alleles observed there (see **Appendix G**), the software computes similarity coefficient between all pairs of spores (see **Appendix B**) and constructs a graph of spores such that two spores represented by nodes are connected with an edge if their similarity coefficient is above the threshold SIMILARITY_COEFFICIENT (the default value is 0.2). Note that this threshold can be adjusted by the user for better performance (see **Appendix C**). Given the graph, the software detects clusters of highly connected spores using the fast algorithm for community detection in large graphs (Clauset et al. 2004). We used the implementation of the algorithm from the python *igraph* library (function community_fastgreedy).

If the user chooses to use clustering of the spores, then the software continues by executing the heuristic search component (steps 1-13 in red in Figure S4). During the heuristic search the software attempts to detect all tetrads within each cluster either by searching for the tetrads directly (step 1) or indirectly using a shadow approach by first computing all triplets (step 3) and then looking for tetrads as a combination of a triplet with another spore from the cluster (step 4). After that, the heuristic search attempts to use the remaining triplets detected within each cluster to detect the tetrads outside the clusters by combining each triplet with each spore outside a cluster – a version of a shadow search (steps 6, 9, and 12). The tetrads, detected at each step of the heuristic search, are removed from the downstream analysis, which considerably reduces the size of the search at later steps.

Once the heuristic search is complete, the software proceeds to execute the direct search component (steps 14-22 in blue in Figure S4). The search takes all the remaining spores and uses the shadow approach to first find candidate triplets (step 14) and then to detect the tetrads that consist of a triplet and another spore. Once the shadow search is complete, the software proceeds to search for tetrads exhaustively (step 17). After that, the software extracts the remaining unused triplets, consisting of three sister spores, and searches for pairs of sister spores among the remaining group.

The last component of the software (step D in yellow in Figure S4) assembles all the detected tuples, labels them, and outputs in a user-friendly format.

## Appendix E

**Figure S6.**
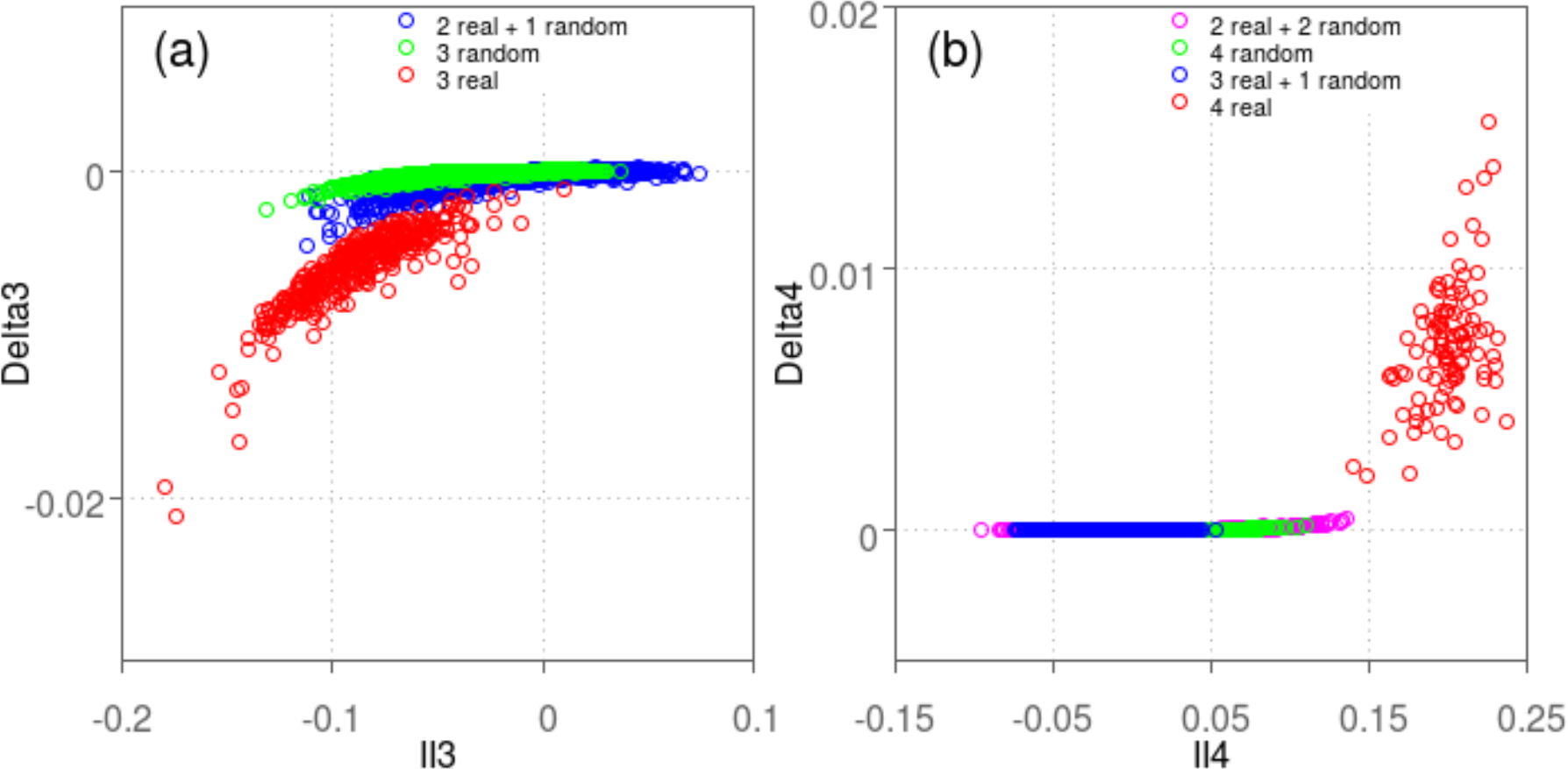
Comparison of delta with interaction information for 3- and 4-spore cases. All measures were computed on the simulated data (1000 markers, 100 tetrads, 1% noise, and 5% missing data). Panel (a) shows the scatter plot of Interaction Information scores versus delta scores computed on all possible groups of 3 spores. Each group is colored red if all 3 spores of the group came from the same tetrad, blue if only 2 spores came from the same tetrad, and green if all spores came from different tetrads. Panel (b) shows the scatter plot of the scores computed on all possible groups of 4 spores. Note that only 1 million randomly sampled tuples are shown for the groups in green (all spores are from different tetrads). [note reversal of sign for II]

**Figure S7.**
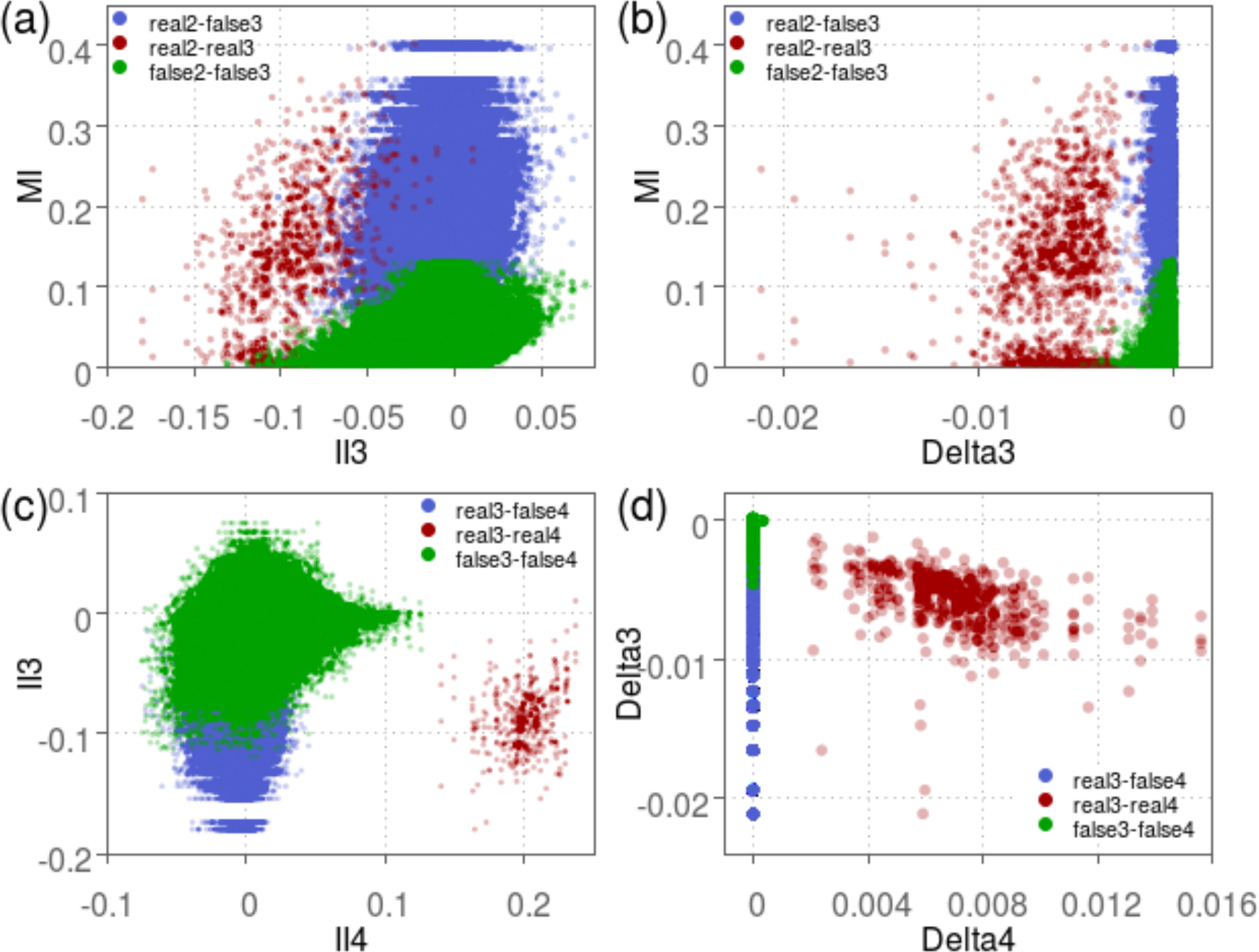
Comparison of the amount of information between 2-spore and 3-spore levels (a-b) and between 3-spore and 4-spore levels (c-d) as measured by interaction information (a, c) and delta (b, d). All measures were computed on the same data as in Figure 2. Panels (a-b) show the scatter plot of scores computed on all possible groups of 3 spores and their two-spore subsets while panels (c-d) show the scores for groups of 4 spores and their 3-spore subsets. Each group is colored red if all spores of the group came from the same tetrad, blue if all spores except one came from the same tetrad, and green otherwise. The scores of the blue and red sets are plotted in their entirety, whereas for the green sets we randomly selected 1 million groups.

## Appendix F

Error-free budding yeast tetrad genotypes were simulated in the form of a table using a custom R script (see *tetrad_sim_1_commented.R* file in the software package), with each row representing a single spore and each column a randomly generated marker position. The number of marker positions and tetrads are specified by the user. Tetrads are encoded as consecutive groups of 4 spores with the first 2 spore rows having the same centromere alleles and the second 2 having the mirror pattern. Parental alleles are encoded as “1” and “2”. In every tetrad, each chromosome experienced one randomly placed obligatory crossover, plus an average (Poisson) of 6 further randomly placed crossovers per megabase (Mancera et al, 2008).

## Appendix G

We estimated the probability that each centromere was derived from the “A” or “B” haplotypes based on the alleles observed at the markers flanking each centromere and the probability of crossovers occurring in the centromere-marker intervals. These probabilities were calculated from the physical sizes of the intervals (in basepairs) using a global estimate of the relationship between physical and genetic distance. This estimate (*b*) was derived by fitting Haldane’s formula with physical distance replacing genetic distance to the observed recombinant fraction between all markers within 250kb of one another. Specifically the formula

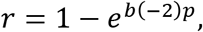

where *r* is a recombinant fraction, *p* is physical distance in bp, and *b* is the conversion factor between genetic and physical distance, was fit using the nls (non-linear least squares) command in R.

As an alternative to this approach, the estimate of total crossovers per meiosis from (Mancera et al, 2008) can be used to provide a coefficient for converting physical to genetic distances in *S. cerevisiae*.

## Appendix H

**Table S1.**
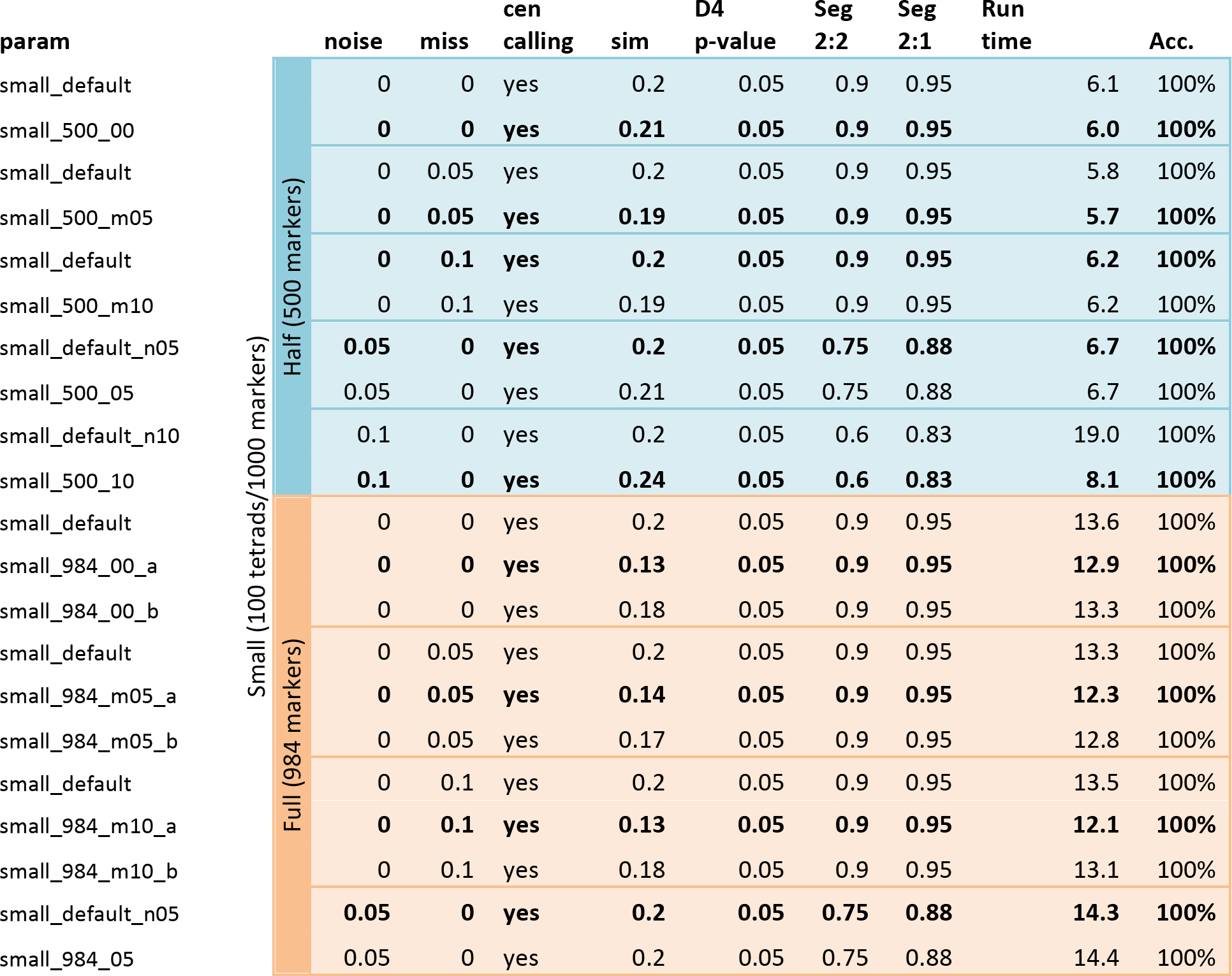
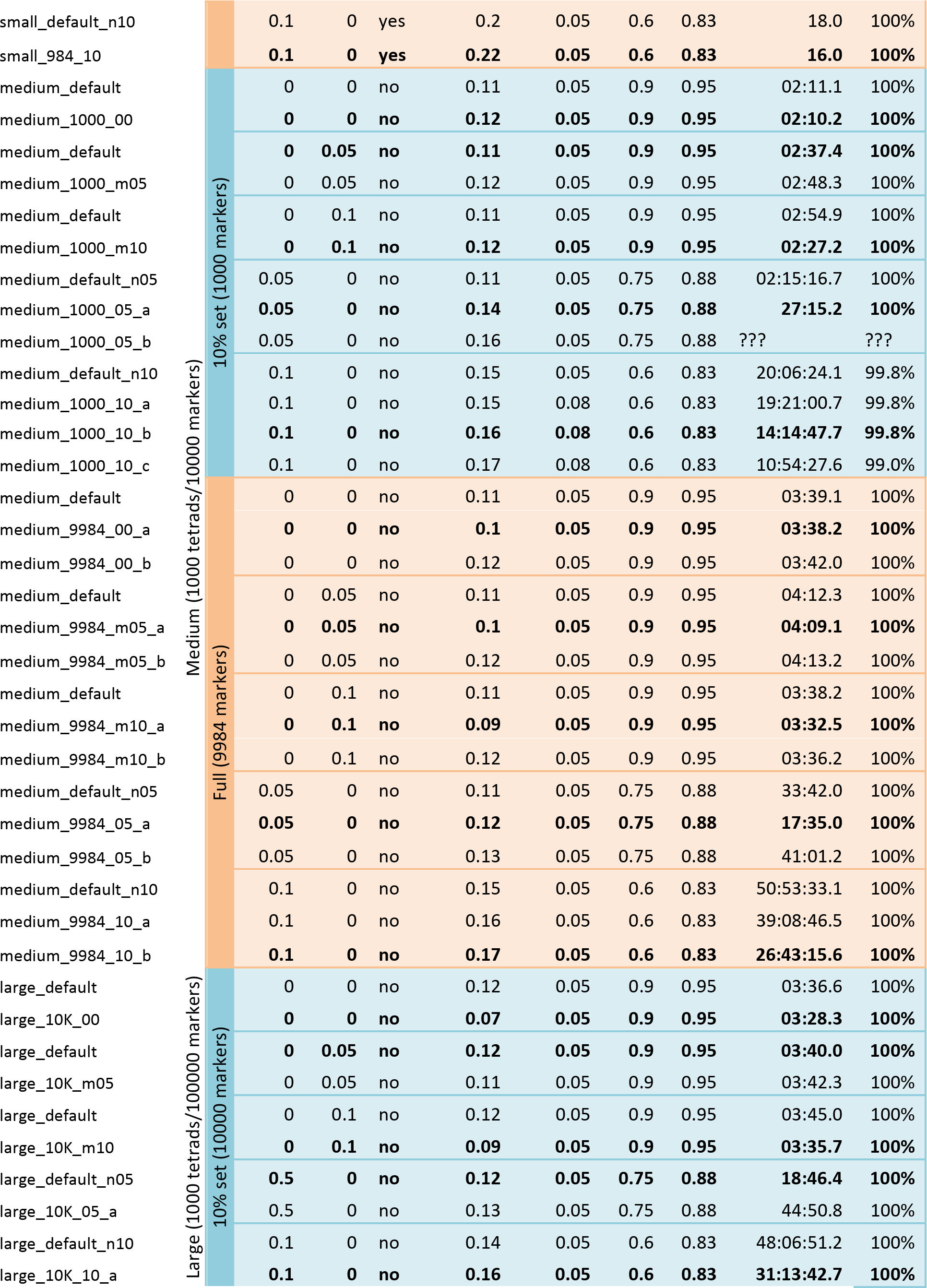
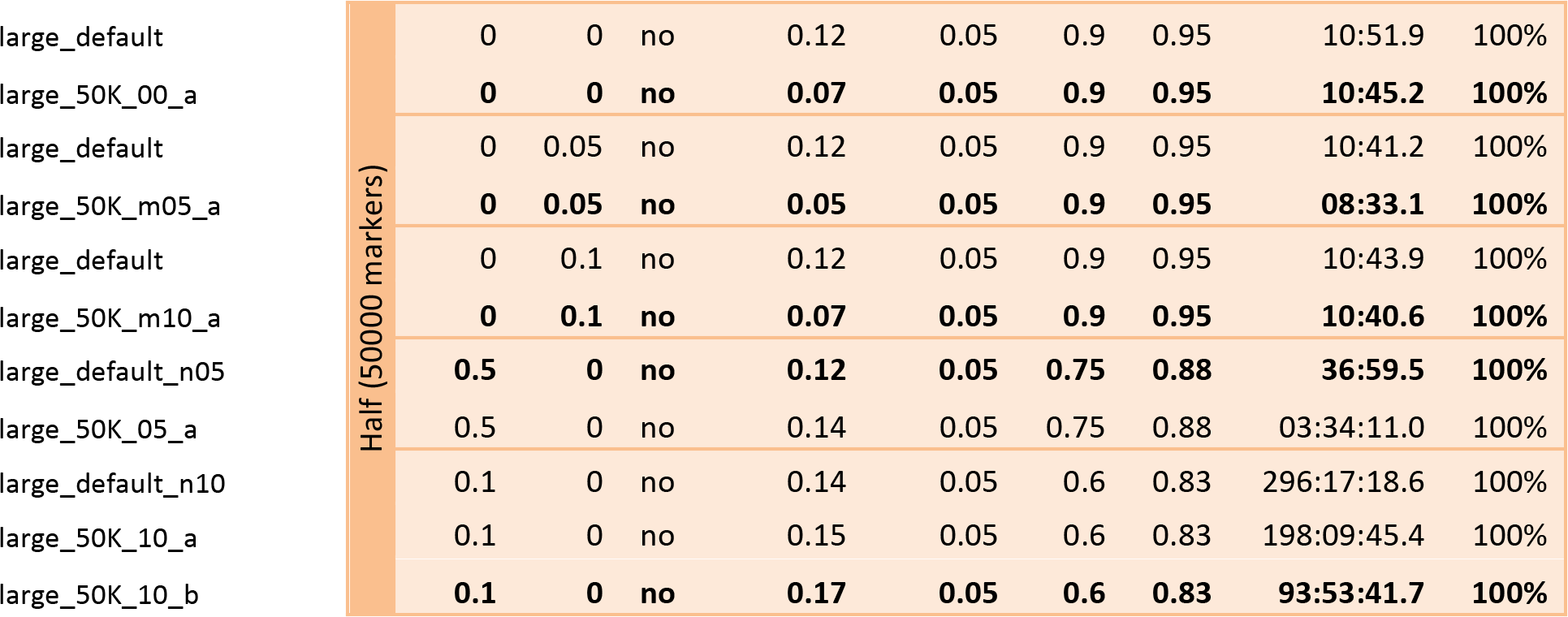
The tetrad detection software applied to various simulated test sets using multiple different parameter settings.

